# Characterization and Engineering of the Type 3 Secretion System Needle Monomer from *Salmonella* Through the Construction and Screening of a Comprehensive Mutagenesis Library

**DOI:** 10.1101/2024.05.02.592225

**Authors:** Lisa Ann Burdette, Samuel Alexander Leach, Nolan Kennedy, Bon C. Ikwuagwu, Jordan S. Summers, Danielle Tullman-Ercek

## Abstract

Protein production strategies in bacteria are often limited due to the need for cell lysis and complicated purification schemes. To avoid these challenges, researchers have developed bacterial strains capable of secreting heterologous protein products outside the cell, but secretion titers often remain too low for commercial applicability. Improved understanding of the link between secretion system structure and its secretory abilities can help overcome the barrier to engineering higher secretion titers. Here we investigated this link with the PrgI protein, the monomer of the secretory channel of the Type 3 Secretion System (T3SS) of *Salmonella enterica*. Despite detailed knowledge of the PrgI needle’s assembly and structure, little is known about how its structure influences its secretory capabilities. To study this, we recently constructed a comprehensive codon mutagenesis library of the PrgI protein utilizing a novel one pot recombineering approach. We then screened this library for functional T3SS assembly and secretion titer by measuring the secretion of alkaline phosphatase using a high-throughput activity assay. This allowed us to construct a first-of-its-kind secretion fitness landscape (SFL) to characterize the PrgI needle’s mutability at each position as well as the mutations which lead to enhanced T3SS secretion. We discovered new design rules for building a functional T3SS as well as identified hypersecreting mutants. This work can be used to increase understanding of the T3SS’s assembly and identify further targets for engineering. This work also provides a blueprint for future efforts to engineer other complex protein assemblies through the construction of fitness landscapes.

**Importance:** Protein secretion offers a simplified alternative method for protein purification from bacterial hosts. However, the current state-of-the-art methods for protein secretion in bacteria are still hindered by low yields relative to traditional protein purification strategies. Engineers are now seeking strategies to enhance protein secretion titers from bacterial hosts, often through genetic manipulations. In this study, we demonstrate that protein engineering strategies focused on altering the secretion apparatus can be a fruitful avenue toward this goal. Specifically, this study focuses on how changes to the PrgI needle protein from the type 3 secretion system from *Salmonella enterica* can impact secretion titer. We demonstrate that this complex is amenable to comprehensive mutagenesis studies and that this can yield both PrgI variants with increased secretory capabilities and insight into the normal functioning of the type 3 secretion system.

## Introduction

Bacteria are popular hosts for recombinant protein production because they are genetically tractable, robust, and inexpensive to culture. Traditionally, bacterial protein expression methods are intracellular, however, and downstream processing requires lysis and recovery of the protein product from a biochemically similar milieu. Recovery of proteins expressed intracellularly in bacteria is often further complicated by low soluble yields caused by host toxicity or accumulation of the protein product in insoluble aggregates^1,2^. Secreting the protein out of the cell has the potential to alleviate these problems while retaining the benefits of bacterial protein production. At least five types of bacterial secretion systems have been shown to secrete recombinant proteins outside the cell, though commercial feasibility remains out of reach due to limitations on titer, substrate compatibility, and efficiency^3–6^.

The type 3 secretion system (T3SS) transports proteins directly from the cytoplasm to the extracellular space and is thus a promising platform for protein secretion in bacteria. It is not required for cellular viability, which allows it to be co-opted solely for heterologous protein production. The T3SS is capable of secreting a wide variety of recombinant proteins^7–9^. However, the system remains unoptimized for secreting products at industrially viable titers, and the engineering space remains largely unexplored. The structure of the T3SS is well-studied and plays an integral part in its function (**Fig 1**). However, little is known about how the structure of the T3SS influences its secretory capability. Learning how to manipulate the apparatus to improve secretion efficiency for recombinant proteins is key to developing the T3SS as a platform for protein production.

**Figure 1.**
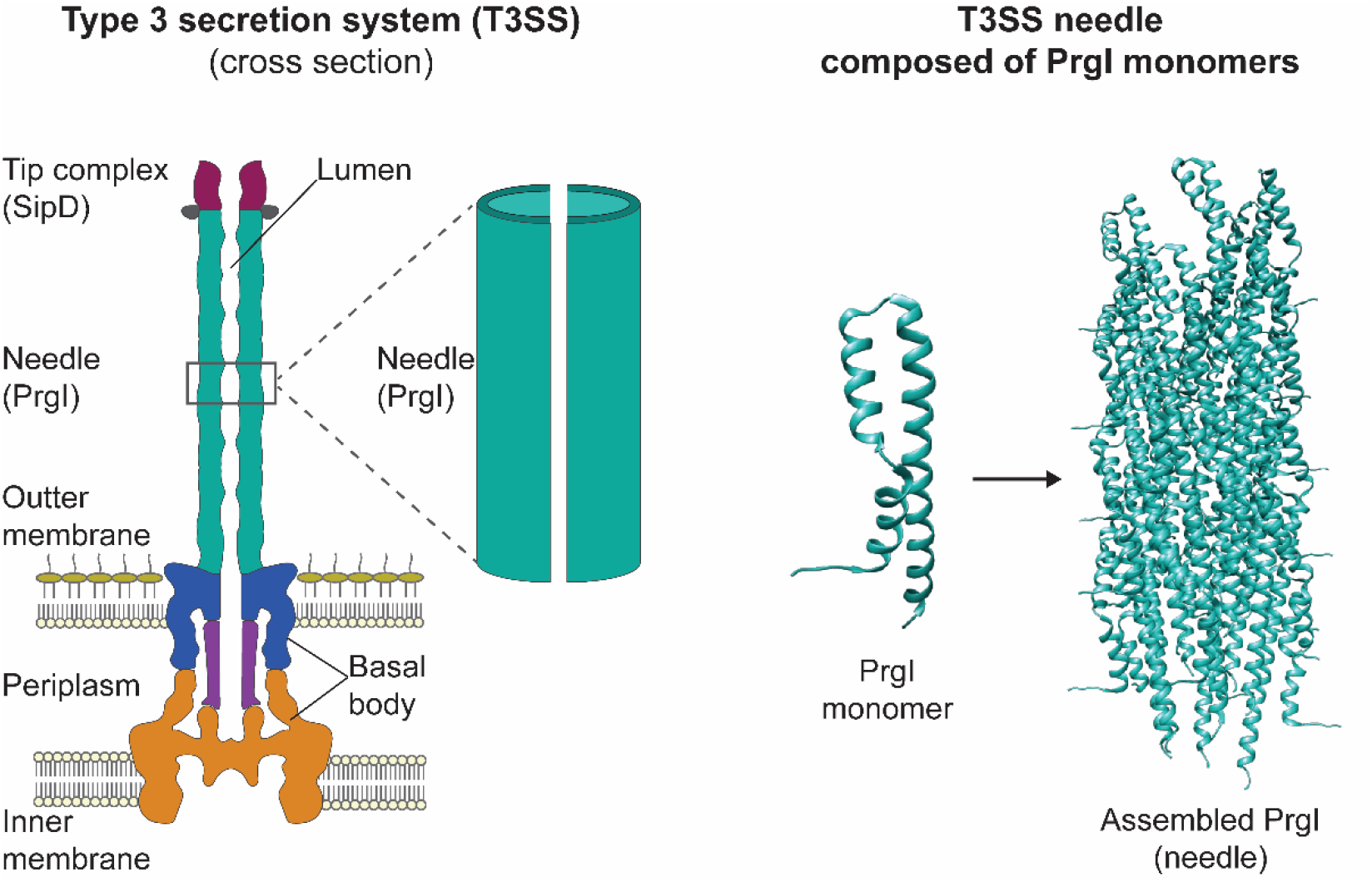
The type 3 secretion system needle is composed of assembled PrgI monomers. (Left) Schematic representation of the type 3 secretion system, which is a large protein complex composed of a basal body that spans the inner and outer cell membranes as well as the periplasm^10^. The needle (teal), composed of assembled PrgI monomers (right) that extend into the extracellular space. The needle is capped by a tip complex composed of the SipD protein. Proteins are secreted from the inside of the cell to the extracellular space through the lumen in the needle complex. PrgI monomers are from the PDB structure 6dwb^11^.

The T3SS is a large (3.5 MDa) protein complex that spans the inner and outer membranes of the bacterial cell^12,13^ (**Fig 1**). A series of rings embedded in the inner and outer membranes comprise the basal body (**Fig 1**). An oligomeric tunnel, rooted in the basal body and composed of about a hundred copies of a single protein, PrgI, extends into the extracellular space (**Fig 1**). The tunnel, or needle, is capped by a tip complex (**Fig 1**). A detailed mechanism of protein translocation through the needle lumen remains unknown, but variations in the proteins that compose the T3SS apparatus, including PrgI, affect its native secretion function^14–17^.

We sought to probe an under-explored area: the impact of variations in T3SS apparatus proteins on secretion. To do so, we chose to use deep mutational scanning and protein fitness landscapes, which are powerful tools for defining protein structure-function relationships and identifying candidates with improved characteristics^18^. This technique is frequently applied to monomeric or small protein complexes with a direct functional output that is readily assayed. Establishing a fitness landscape for self-assembling proteins can be more challenging, as any functional screen or selection reports on both assembly and the functional output.

We used this combined output to our advantage, however, and adapted the recently developed SyMAPS method^19^ to produce a “secretion fitness landscape” (SFL) of the *Salmonella enterica* Typhimurium T3SS needle protein PrgI. Briefly, we created a first-of-its-kind genomically-integrated comprehensive codon mutagenesis (CCM) library and ranked the library members by secreted protein titer. Next generation sequencing of the ranked variants enabled construction of an SFL. The SFL revealed design rules for a functional T3SS needle and provides a blueprint for future engineering targets for increased secretion titer. Through this process, we also were able to identify and confirm new hypersecreting variants. Beyond this immediate application, the work provides a blueprint for the use of comprehensive library design and high-throughput screening to characterize and engineer other multimeric protein structures, including other T3SS proteins as well as the components of other secretion systems.

## Results

### Validating the construction and screening of a genomically-encoded PrgI variant library

PrgI and several of its homologs have been analyzed through alanine scanning and targeted substitution^14,15,20,21^. Single amino acid changes produced variable secretion phenotypes, so we hypothesized that a combined CCM and library screening approach could reveal hypersecreting variants of PrgI. To accomplish this, we validated a genomic library construction and screening strategy. SPI-1 T3SS assembly and activation is a tightly controlled, highly orchestrated process^12,22^, and SPI-1 genes have overlapping regulatory elements, particularly in the *prg* operon where *prgI* is located^23,24^. Thus, we sought to construct a genomically-encoded library to study only the effect of structural changes in PrgI by maintaining the native regulatory structure and induction cascade.

We chose to employ λ Red recombineering^25^ to create the library because it enables construction in a single step. To our knowledge, this technique has not been used in library construction before we carried out this study, so we performed a pilot test by creating a saturation mutagenesis library at only PrgI position 41 to validate both the library construction and screening methods (**Fig 2A**). PrgI^P41^ was of interest because an alanine substitution at the homologous position in MxiH, the needle protein for the *Shigella flexneri* T3SS injectosome, increased secretion of native substrates^15^. The template strain for library construction contained a copy of alkaline phosphatase fused to the T3SS secretion tag SptP integrated at the *sptP* locus, providing a reporter for secretion titer through use of an enzyme activity assay (**Fig 2A**)^26^. It also contained selectable markers at the *prgI* locus, to be replaced by the *prgI* variants (**Fig 2A**). An equimolar pool of all 20 fragments of the saturation mutagenesis library was introduced into this *sptP::sptP*^1^*^-^_167_-phoA prgI::catG-sacB* strain using a single λ Red recombineering event, and 68 of the resulting clones were Sanger sequenced (**Fig 2A**). Of those 68 clones, 88% contained *prgI* alleles that successfully replaced the *catG-sacB* cassette, and all variants were present except PrgI^P41C^ (**Supplemental Fig 1**).

**Figure 2.**
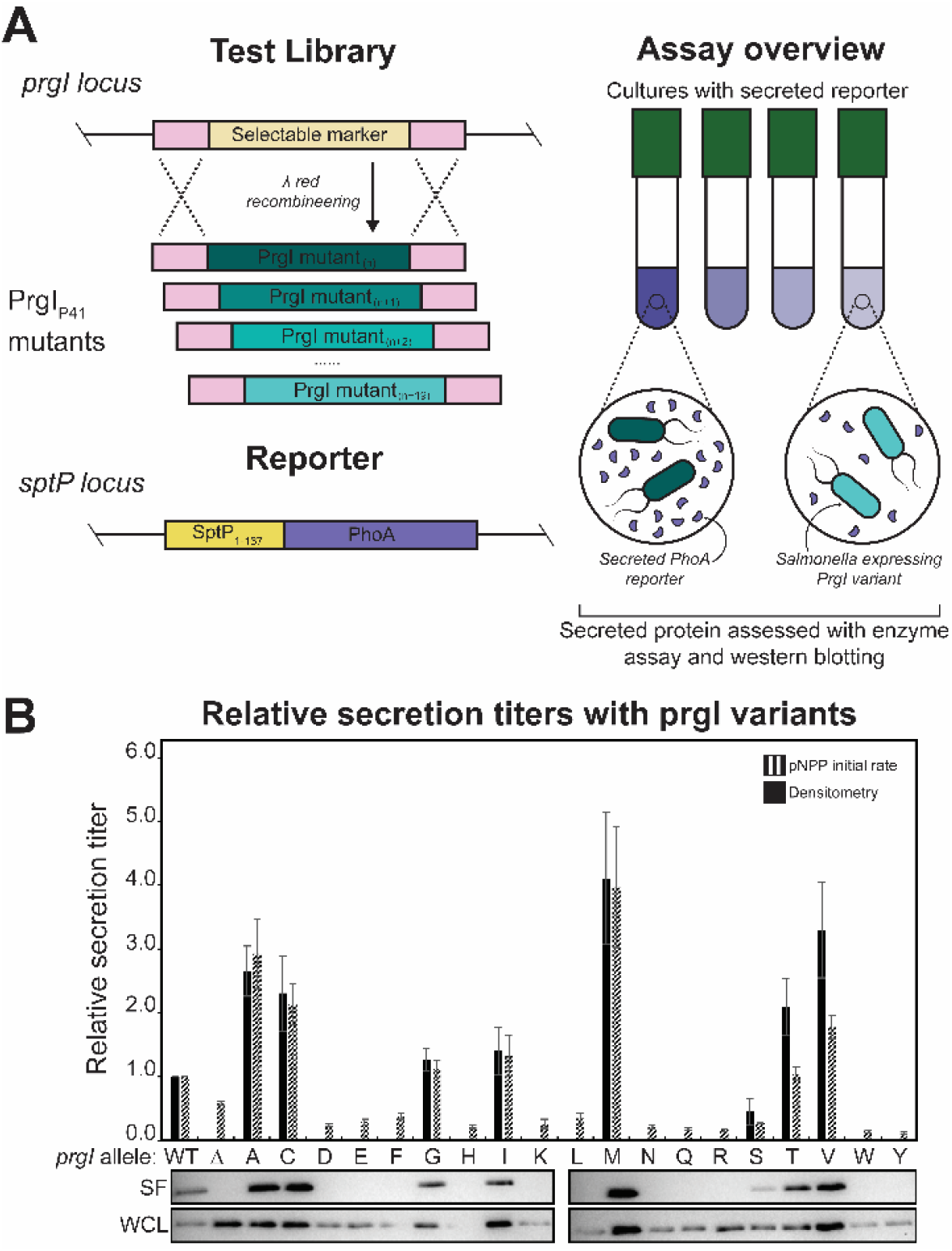
PrgI variants can be screened for activity. (A) Schematic of the library generation and screening methods for assessing PrgI variants. Genes encoding PrgI variants were cloned into the *prgI* locus in a strain containing a SptP-tagged PhoA (alkaline phosphatase, “AP”) reporter at the *sptP* locus. Variants were then assayed for enzyme activity as a proxy for secretion titer. (B) SptP-AP-2xFLAG-6xHis was secreted from ASTE13 *sptP::sptP(1-167)-phoA-2xFLAG-6xHis* and the specified PrgI mutants in LB-L. “Δ” is ASTE13 *prgI::catsacB*. Relative secretion titer was measured using densitometry from western blots and normalized to a PrgI^WT^ strain. Western blots are representative of four biological replicates. Relative secretion titer was also measured by averaging the slope of absorbance at 405 nm versus time for four biological replicates and normalizing to PrgI^WT^. Absorbance at 405 nm was recorded every 5 minutes for 16 hours on a BioTek Synergy HTX plate reader at 37°C. Error bars represent standard error. Representative western blots of the secreted fraction (SF) and the whole culture lysate (WCL) are shown below.

The PrgI^P41^ saturation mutagenesis library produced variable secretion titers, and several variants increased secretion titer compared to wild-type PrgI (PrgI^WT^) (**Fig 2B**). We used the initial rate of secreted alkaline phosphatase activity as a proxy for secreted enzyme titer^27^, and all variants were normalized to PrgI^WT^ (**Fig 2**). Relative secretion titers as assessed by alkaline phosphatase activity generally agreed with western blotting results (**Fig 2B**). The high rate of substitution at the PrgI locus and the high coverage of the PrgI^P41^ library gave us confidence that we could construct and screen a full PrgI library, and the variable secretion titers convinced us of the utility of such a screen.

### Generating a secretion fitness landscape of PrgI

With the genomically-encoded library construction and screening methods validated, we constructed the full CCM library of PrgI. We created the library in a single λ Red recombineering event using the *sptP::sptP*^1–167^–*phoA prgI::catG-sacB* strain and an equimolar pool of synthetic, double-stranded *prgI* gene fragments, each containing a single amino acid change (**Fig 3**). The first and last six amino acids were not changed because they are essential for assembly ^15,20^ and wild-type (WT) residues were not present in the library. Because secretion titer is a phenotype necessarily separate from genotype (the secreted protein is physically separated from the cell and its associated genetic information), clones were screened individually in 96-well plates (**Fig 3**). Glycerol stocks were maintained in the same array as the samples measured for secretion titer to provide the link between genotype and secretion titer (**Fig 3**). We screened 4406 clones to capture threefold the variant library size with a cushion for false positives from the recombineering process. Secretion titer, as measured by the initial rate of secreted alkaline phosphatase activity, was normalized to that of PrgI^WT^ to allow comparison across assay plates. The unscreened library contained 99% of expected variants, and the screened clones captured 90% of the library. After the initial screen, clones were re-arrayed according to relative secretion titer and sorted into ten pools for high-throughput sequencing (**Fig 3**).

**Figure 3.**
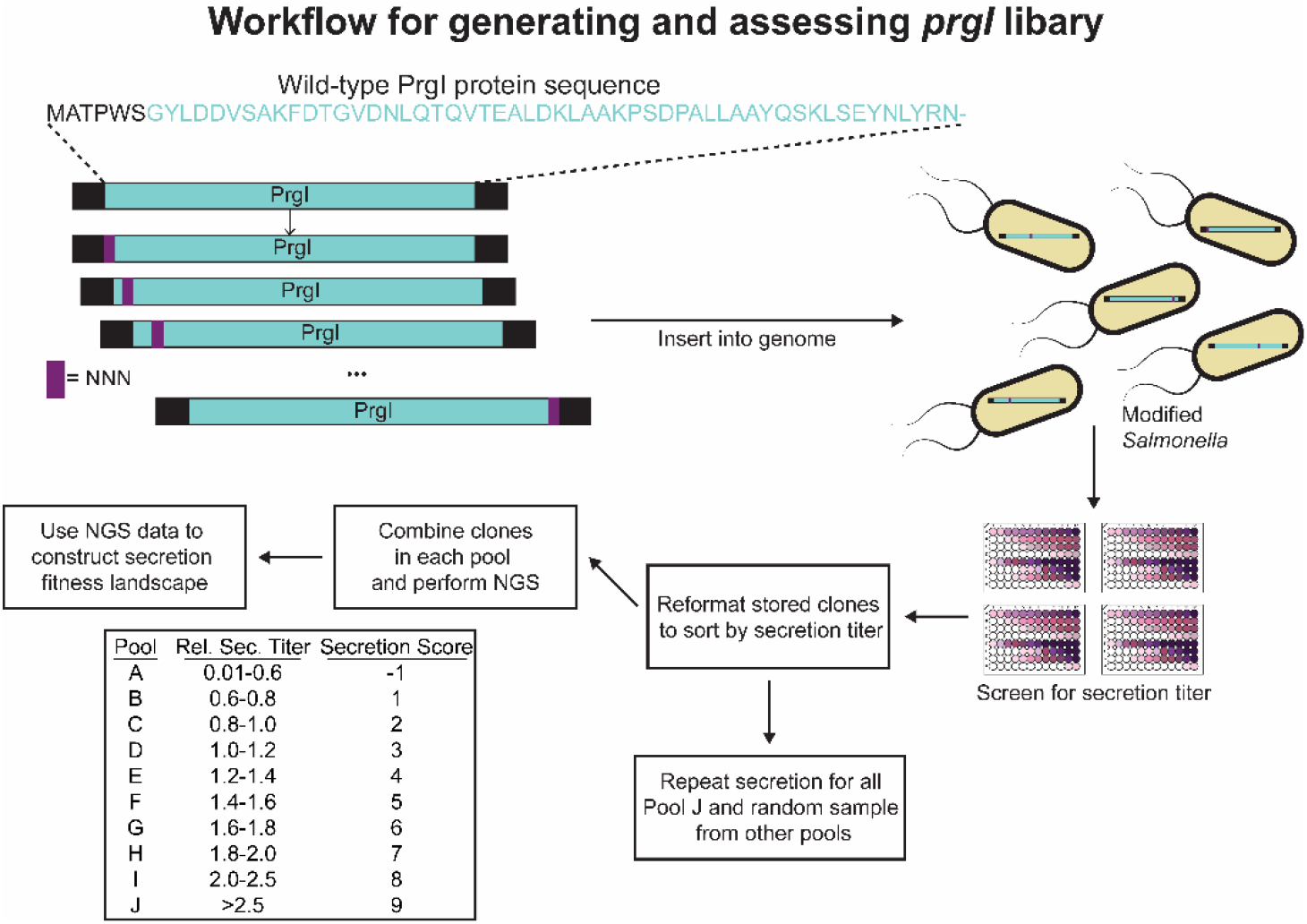
Workflow for PrgI library assembly, screening, and analysis. Gene blocks coding for a single amino acid change in PrgI were introduced to ASTE13 *sptP::sptP*(1–167)-*phoA-2xFLAG-6xHis prgI::catsacB* as a mixture in a single recombineering event to create the library. Individual colonies were inoculated for secretion and screened for alkaline phosphatase activity as described in Methods. The randomly arrayed clones were sorted according to relative secretion titer, combined into their assigned pools, and prepared for sequencing on an Illumina MiSeq.

Dividing clones into pools according to relative secretion titer allowed assignment of a “secretion score” to each variant (**Fig 3**). If variants appeared in multiple pools, a weighted-average secretion score was calculated using the relative abundance of the variant in each pool as weights. A window of likely pool appearances was defined by the alkaline phosphatase activity assay coefficient of variation (CV), and scores outside of that window were not included in the weighted average calculation (see **Supplementary Methods**). This scoring led to construction of a quantitative “secretion fitness landscape” (SFL) that reports on both functional substitutions and those that increased secretion titer (**Fig 4**). WT secretion levels are expected to fall within scores 2-5, variants with scores less than 1 are expected to be non-functional, and variants with scores above 7 are anticipated to confer secretion titers higher than PrgI^WT^.

**Figure 4.**
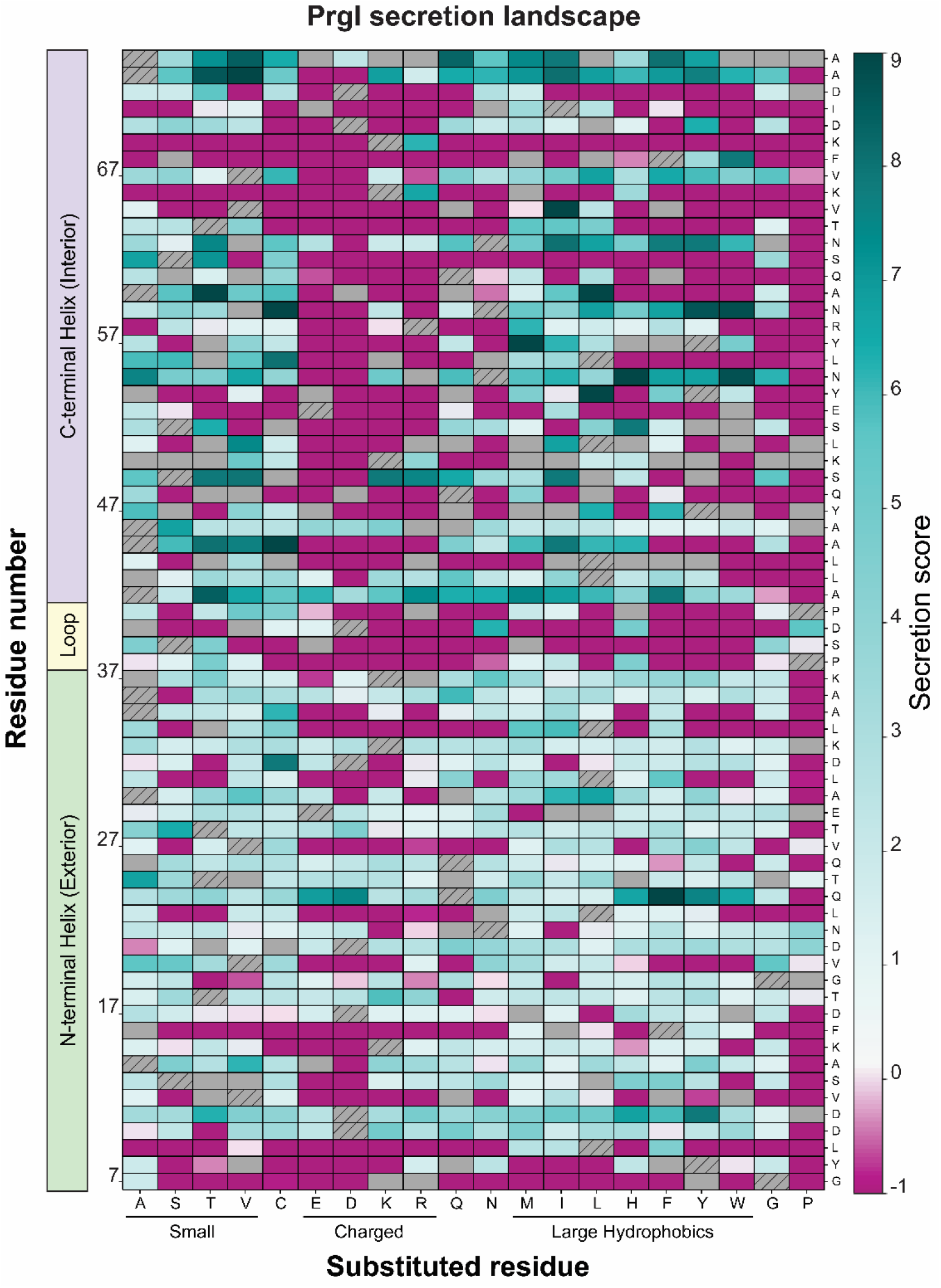
Weighted-average secretion scores for all single amino acid variants of PrgI. The PrgI library was constructed and screened for relative secretion titer using the alkaline phosphatase (AP) activity assay. Variants were sorted into ten pools according to relative secretion titer and sequenced. If variants appeared in multiple pools, a weighted average secretion score was used. Wild-type residues not present in the library are indicated by hatches. Grey boxes denote variants that did not appear in any sequenced pools; i.e., those variants were not screened. Dark teal indicates higher secretion titer while dark pink indicates no secretion.

### The SFL is experimentally validated and reflects known structural constraints

The synthesized library did not contain WT residues or stop codons, nor was any amino acid encoded with synonymous codons, so there were no controls within the SFL to assess validity of the method internally. Instead, we validated the SFL experimentally. All clones from the highest secreting pool (Pool J) and a random selection of 30 additional clones distributed across the remaining pools (Pools A-I) were Sanger sequenced and re-evaluated individually in triplicate using initial rate of secreted alkaline phosphatase activity as a proxy for secretion titer (**Fig 5**). Secretion titer increased with secretion score (R^2^ = 0.87), and all variants in Pool J showed secretion titers at least 50% greater than PrgI^WT^ (**Fig 5**). Combined, these results confirm the ability of the secretion score to predict general secretion behavior.

**Figure 5.**
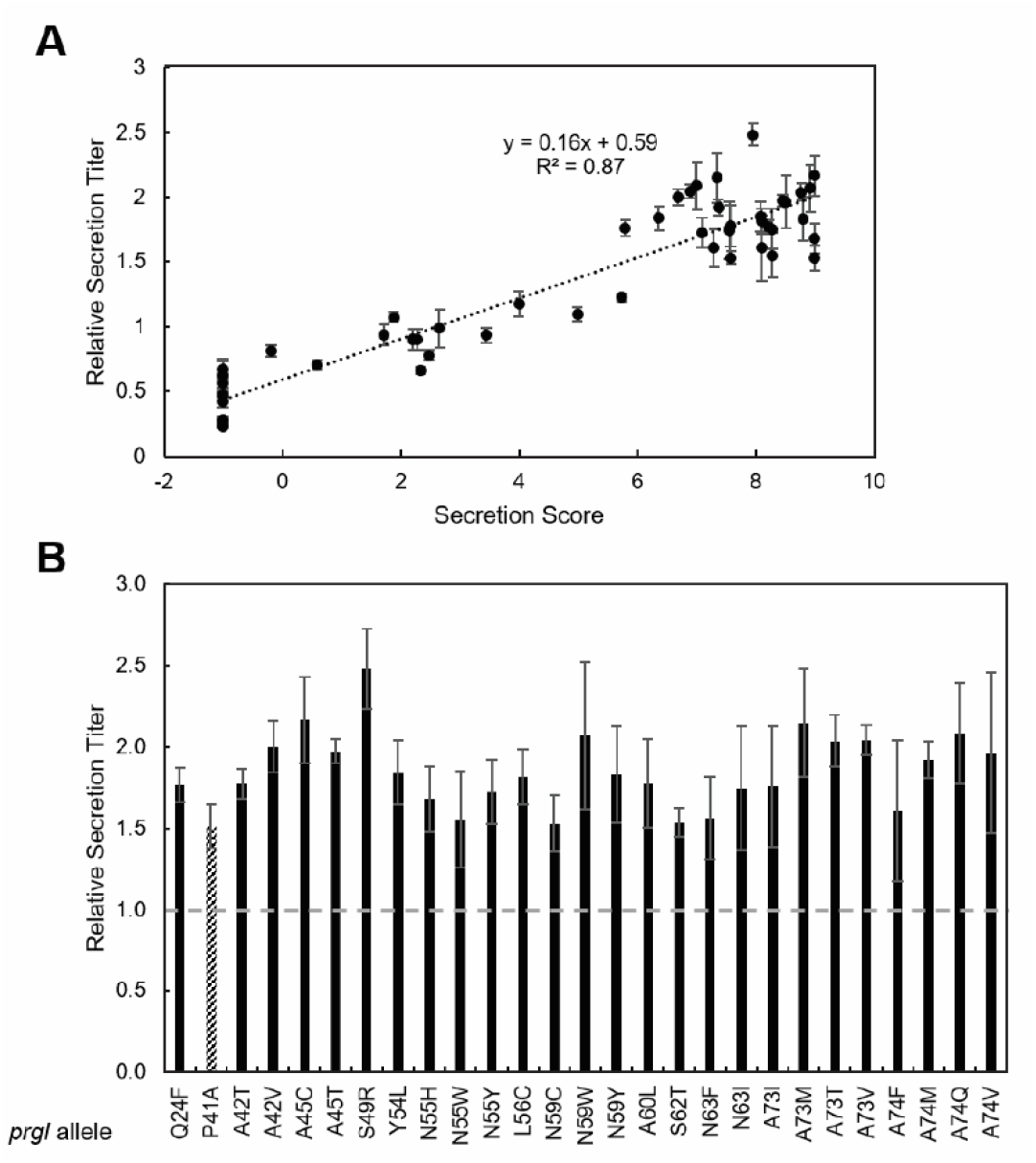
Weighted-average secretion scores predict relative secretion titer. All clones from the highest secreting pool and 30 random clones from the remaining pools were patched onto LB-agar plates from the reformatted glycerol stocks sorted by pool (see supplementary methods for details on reformatting). Patched colonies were inoculated for secretion titer measurement, which followed the same workflow as library screening. Clones were Sanger sequenced, and the secretion score was plotted against the newly measured secretion titer (A). Error bars represent standard error of three biological replicates. Clones from the highest secreting pool were plotted separately to highlight differences (B). WT-level secretion titer is indicated by a dotted line. PrgI^P41A^ (stripes) was included for comparison as the previous best secreting variant. Error bars represent one standard deviation to allow direct comparison among variants.

The SFL can also be validated by examining poorly substituted residues in the context of available structural information. To determine the overall mutability of each position in PrgI, an average secretion score was calculated by averaging the secretion scores of all variants at each position (**Fig 6, right**). **Table 1** lists all residues with an average secretion score < 0.5 i.e. most or all variants were non-functional, making the position immutable. For each of these residues, we calculated conservation and buried surface area (BSA) (**Table 1**, **Fig 6, left and middle**). Conservation is a measure of how similar the amino acid sequence is between protein homologs in different organisms. If a residue is highly conserved, it indicates that the position is likely crucial in the function of the protein and cannot be mutated. We calculated conservation from all members of Pfam group PF09392, which are T3SS needle homologs across enteric bacteria. BSA is a measure of how accessible a residue is to solvent in the context of the protein complex. Residues with a high BSA are likely crucial to the protein’s structure and cannot be mutated without affecting protein folding or structure. PDBePISA yielded BSA for each residue and each monomer in PDB 6dwb^28^, and BSA was averaged across all monomers and normalized to the averaged accessible surface area per residue. For the poorly substituted residues listed in **Table 1**, most were buried within the structure, conserved, or both (**Fig 6**), which suggests that the native amino acid at those positions plays an important role in the needle structure. Indeed, recently solved structures of the needle show that all residues in **Table 1** participate in inter- and intra-subunit interaction networks that stabilize the needle filament or are essential for secretion activity^17,29,30^.

**Figure 6.**
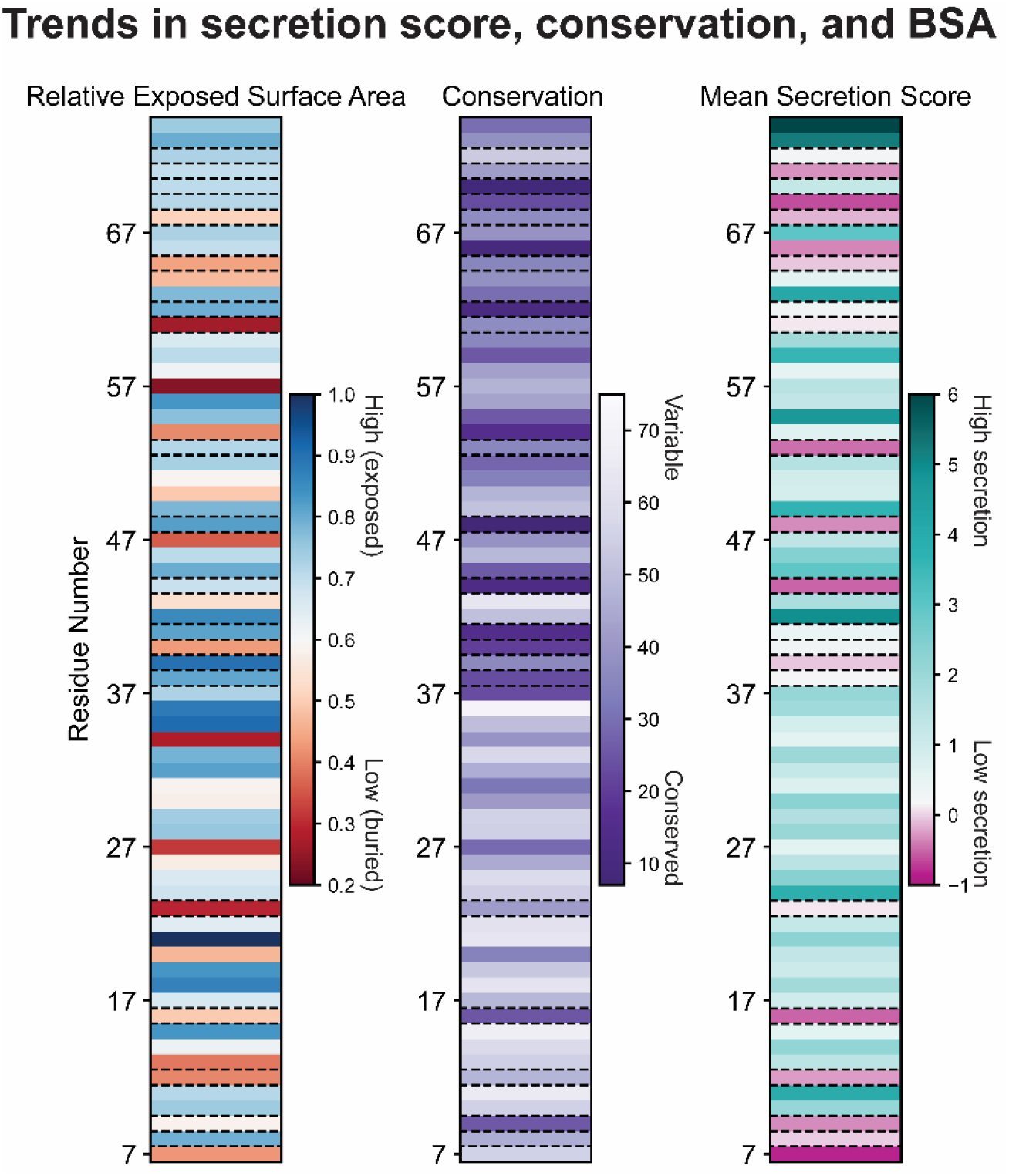
Trends in secretion score, conservation, and buried surface area for PrgI variants. Relative exposed surface area of PrgI residues (high in blue, low in red) (left), relative conservation of PrgI residues (conserved in purple, variable in white) (middle), and mean secretion score for PrgI variants (high secretion in teal, low secretion in pink, WT secretion in white) (right). Immutable residues are highlighted with dashed lines.

**Table 1.** BSA and conservation scores for poorly substituted residues.

| Native Residue | Normalized BSA | Conservation Score |
| --- | --- | --- |
| G7 | 0.51 | 2.42 |
| Y8 | 0.25 | 3.94 |
| L9 | 0.55 | 7.16 |
| V12 | 0.74 | 3.68 |
| F16 | 0.98 | 7.24 |
| L23 | 0.48 | 4.59 |
| P38 | 0.56 | 7.55 |
| S39 | 0.51 | 5.44 |
| D40 | 0.37 | 7.98 |
| P41 | 0.49 | 8.81 |
| L44 | 0.45 | 9.12 |
| Q48 | 0.51 | 10.00 |
| E53 | 0.52 | 5.57 |
| Q61 | 0.72 | 5.29 |
| S62 | 0.53 | 9.48 |
| V65 | 0.87 | 5.59 |
| K66 | 0.28 | 9.62 |
| F68 | 0.54 | 5.30 |
| K69 | 0.25 | 7.52 |
| I71 | 0.56 | 4.53 |
| D72 | 0.48 | 2.72 |

### Amino acid properties reveal substitution preferences

Examining the preference for ten amino acid properties at each position condensed the SFL and revealed substitution preferences (**Fig 7**). To do this, we grouped amino acids according to particular properties including volume, molecular weight, length, steric hindrance, polarity, polar area, fraction water, hydrophobicity, nonpolar area, and flexibility (**Fig 7**). We averaged the secretion scores of these amino acid substitutions at each position to calculate a property secretion score. Substitutions that increased secretion titer were often large and hydrophobic (**Fig 7**). The residues that accepted few functional substitutions, i.e. those with low average secretion scores, preferred no amino acid property (**Fig 7**). Many residues, especially in the N-terminal helix (residues 7-37), had functional substitutions with all types of amino acids. In general, polar and flexible amino acids were disfavored, often together in a repeating pattern throughout PrgI (see the negative scores for polar substitutions at residues 23, 27, 31, etc). The native amino acids in this pattern were hydrophobic except at positions R58 and T64.

**Figure 7.**
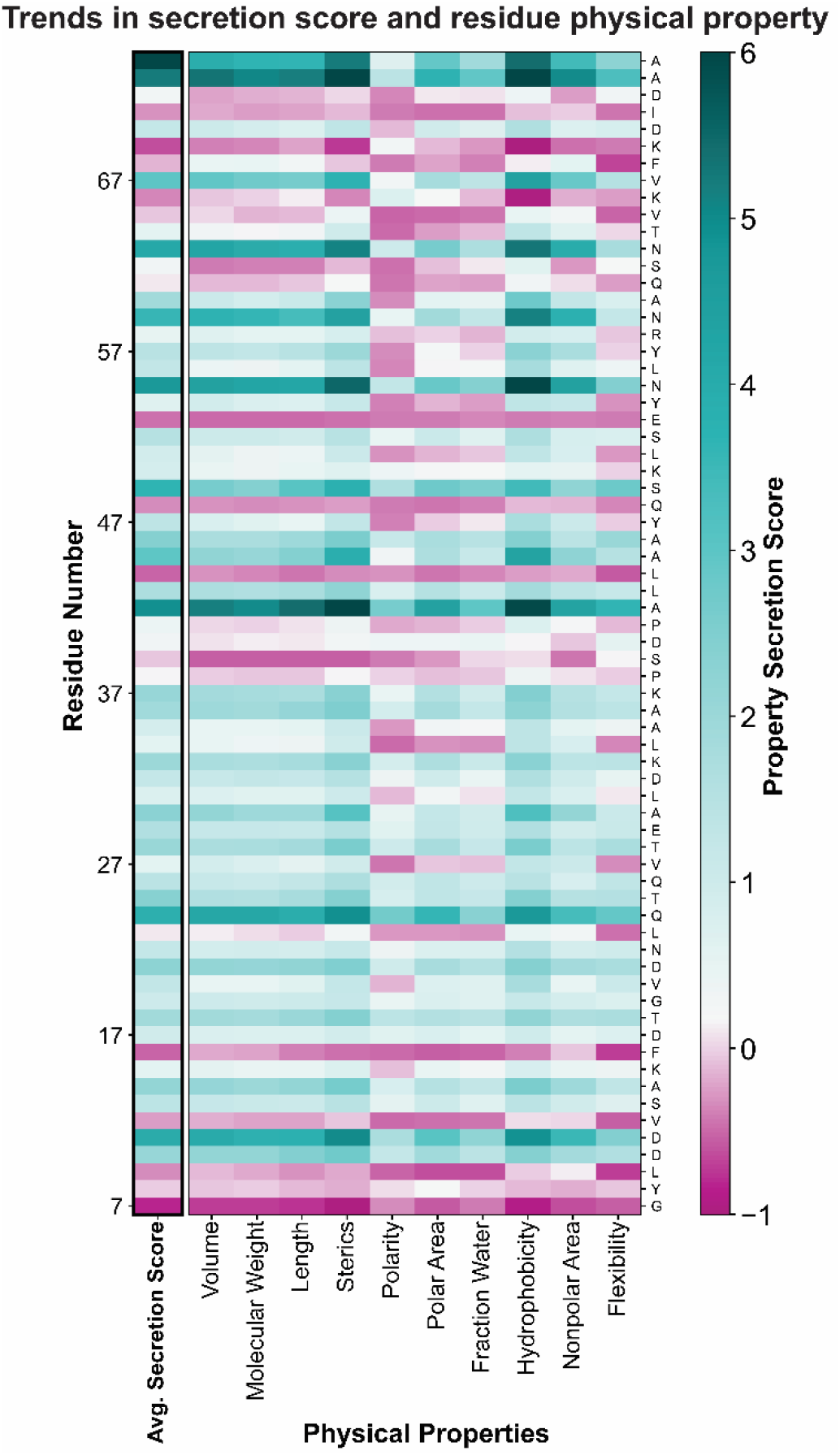
Trends in amino acid property and secretion scores. Amino acids were grouped by the physical properties shown on the x-axis (bottom), and the secretion scores of each group at each position were averaged to calculate a property secretion score. Low secretion is shown in pink, high secretion is shown in teal, WT-like secretion is shown in white.

This preferential property patterning, tendency of hydrophobic residues to increase secretion score, and the observation that buried and conserved residues were generally immutable (**Table 1**, **Fig 6**) led us to hypothesize that conservation and BSA could serve as predictors of mutational tolerance. Additionally, we hypothesized that patterns in mutability would correspond to structural components of PrgI. To test this idea, we calculated conservation and BSA as before for all residues. We clustered positions according to secretion score to better discern patterns in the data. The clustering produced three groups that roughly corresponded to high, medium, and low average secretion scores (**Supplemental Fig 2**). Importantly, residues that were both highly conserved and buried had low mutational tolerance as expected, but that was the limit of the predictive power of conservation and BSA on positional mutation tolerance. Residues with both low BSA and low conservation allowed many functional substitutions, but few increased secretion titer above PrgI^WT^ (**Supplemental Fig 2**). As expected from the prior observation of substitution preferences by property, the highest secreting PrgI variants (cluster in the top left corner of **Supplemental Fig 2**) were often large hydrophobic residues in the C-terminal helix.

### The SFL reveals structural substitution patterns

PrgI is composed of two helices connected by a four-amino acid loop (**Fig 8A**). The N-terminal helix is exposed to the exterior environment, while the C-terminal helix is packed into the structure and forms the needle interior (**Fig 8A**). Mapping the average secretion score per residue onto the needle structure confirmed a pattern suggested by the SFL and clustering (**Fig 4 and Fig 8**) — many substitutions were allowed in the N-terminal helix (**needle exterior, Fig 8B, left**), but few increased secretion titer above PrgI^WT^. Conversely, few substitutions were functional in the C-terminal helix (**needle lumen, Fig 8B, right**), but functional substitutions frequently increased secretion titer residues (**Fig 8B, dark teal substitutions**).

**Figure 8.**
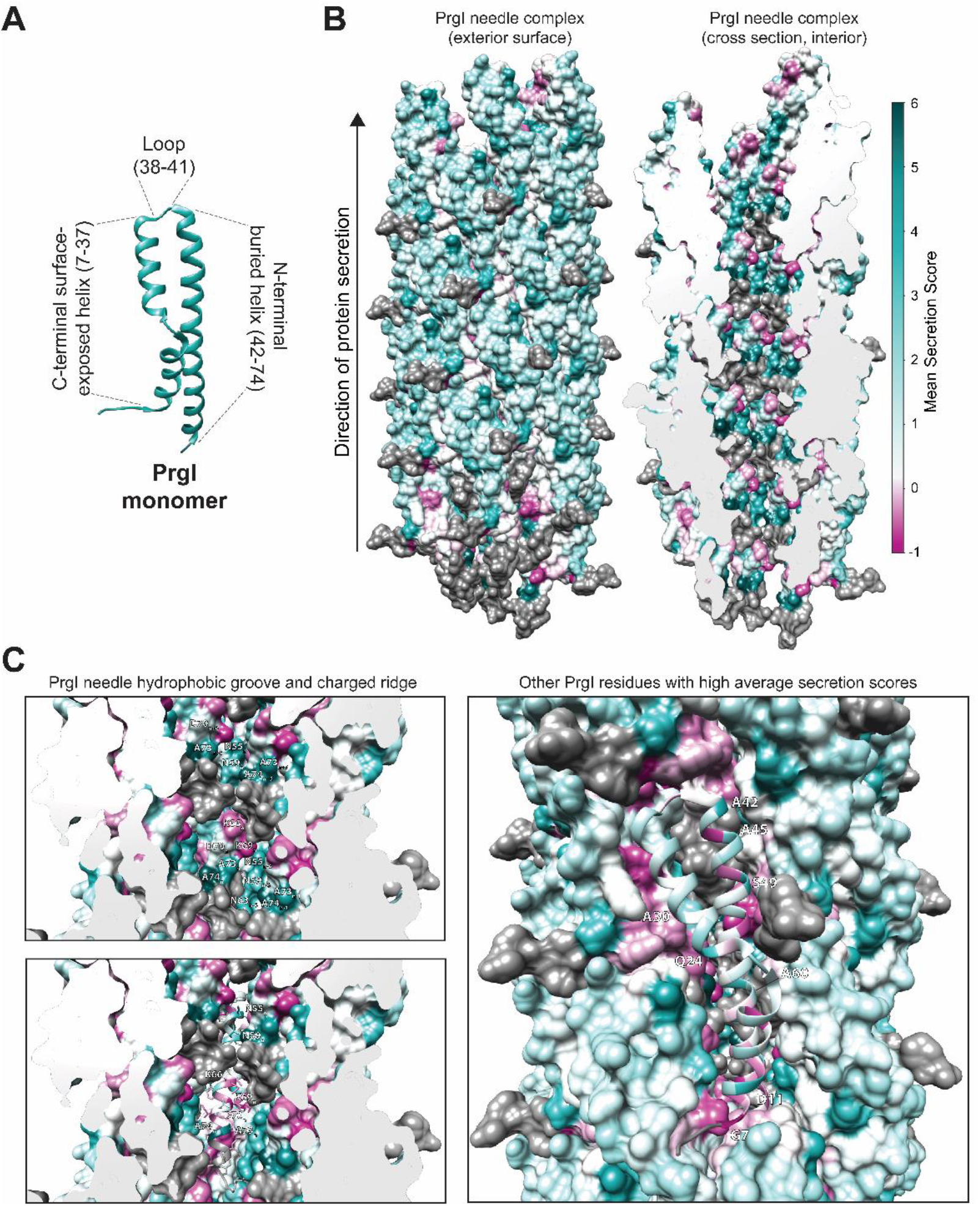
Structural insights from secretion fitness landscapes. (A) The needle and monomer structures are PDB = 6DWB^11^ from Chimera^28^. The C-terminal helix comprises residues 7-37, the loop comprises residues 38-41, and the N-terminal helix comprises residues 42-74. (B) Interior and exterior residues show different patterns for secretion fitness. Secretion scores were averaged across all substitutions at each residue and mapped on to the PrgI needle structure. Dark grey denotes residues that were not modified in the library design. (C) Substitution with large hydrophobic amino acids increased secretion titer at residues that line the hydrophobic groove of the needle interior. The hydrophobic groove in the needle interior is composed of alternating N55, N59, A73, and A74 from the indicated chains. Residues L66, L69, Q77, and R80 form the charged, raised groove. Residue 70 contributes to the charged raised groove with its native aspartic acid but also tolerated several amino acid substitutions. A ribbon colored by average secretion score shows the predicted orientations of each amino acid.

The needle interior shows a stark substitution pattern, with alternating helical bands of well-substituted and poorly substituted residues (**Fig 8B-C**). The band of poorly substituted residues includes those C-terminal residues excluded from the library (**residues in grey, Fig 8**) because they are essential for needle assembly^15,20^. Models of the PrgI needle depict the needle interior as a right-handed groove with alternating charged and hydrophobic residues forming the edges and lumen of the groove, respectively^30^ (**Fig 8C**). The raised, charged groove is highly conserved across all species with a T3SS^29,30^, so we were not surprised to discover that most modifications at those residues disrupted secretion.

Poorly substituted residues also occurred at the interfaces between helices and adjacent monomers (**Fig 8C**). A common theme of poorly substituted residues, aside from degree of burial and conservation, was that the mutations that were tolerated were of similar character to the native residue (**Fig 7**). This supports the hypothesis that these interfacial residues facilitate proper structural arrangement and packing of each monomer within the needle structure.

Of the 63 variants with secretion scores greater than 7, 55 were in the C-terminal helix. We were surprised to discover that half of those 55 favorable substitutions were present at residues 55, 59, 63, 73, and 74, which form or contact the hydrophobic groove in the needle lumen (**Fig 8C**). The hydrophobic groove is fairly well-conserved (**Fig 6**), so it was surprising to find that many mutations were not only tolerated but significantly increased secretion titer at these positions. Further, the most favorable substitutions at those residues were larger and more hydrophobic amino acids (**Fig 4**, **Fig 7**).

Well-substituted residues, or those with high average secretion scores, also included positions 11, 24, 30, 42, 45, 49, and 60 (**Fig 8C**). Substitutions at residues 30, 45, 49, and 60 likely change inter- or intra-monomer interactions, as those residues face neighboring chains (**Fig 8C**). Residues 11, 24, and 42 face the needle exterior (**Fig 8C**), so the beneficial substitutions at those residues must affect another aspect of T3SS assembly, solubility, or expression. Notably, residues 11 and 49 may be important for forming contacts with the tip protein SipD^17^.

### Beneficial mutations are additive with other secretion titer enhancements

Secretion titer via the T3SS is maximized in an optimal growth medium with plasmid-based expression of the secreted protein fused to the secretion tag SptP and overexpression of a T3SS master regulator, *hilA*^5,9^. Thus, we sought to evaluate whether the beneficial PrgI mutations revealed by this study caused a general increase in secretion titer and were additive with those existing strategies. We selected four top clones: A45C, S49R, N59W, and A74Q. N59W and A74Q face the needle interior and have a different character than the native amino acid. A45C and S49R likely affect inter-subunit interactions and/or interactions of PrgI with other T3SS components. In combination with *hilA* overexpression and in an optimized medium, all four variants increased secretion titer of two model proteins, the human domain of intersectin (DH) and recombinant human growth hormone (HGH), at least 50% above PrgI^WT^, indicating that the variants were additive with the other improvements (**Fig 9**). There was no significant difference among the variants, suggesting that a twofold increase in secretion titer was an upper limit for these variants in combination with other enhancements of secretion titer.

**Figure 9.**
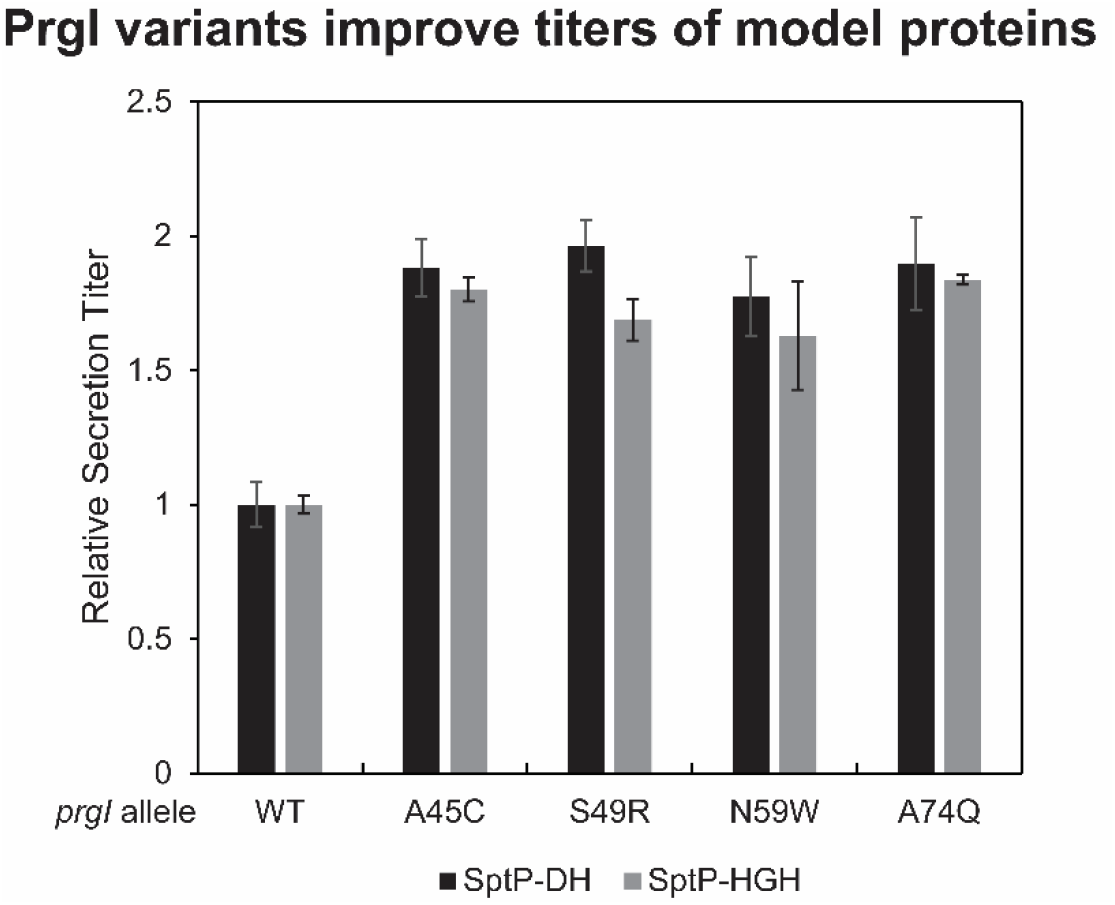
High secreting PrgI variants enable increased secretion of model proteins. The WT strain along with four of the top secreting variants from the PrgI library were transformed with the *hilA* overexpression plasmid and a plasmid expressing either SptP-DH or SptP-HGH. Each strain was grown in an optimal growth medium. Secretion samples were collected after hours, and relative secretion titer was measured via semi-quantitative western blot. Error bars represent the standard error of three biological replicates.

## Discussion

Protein engineering relies on either building large libraries of randomly generated variants or small libraries of rationally designed variants. However, as the cost of DNA synthesis and sequencing has fallen, we are now able to construct CCM libraries, which encompass every possible individual mutation across a protein sequence. These libraries combine the advantages of the breadth in sequence space covered by a random mutagenesis library with the depth of a rationally designed approach. While CCM has been applied to both individual proteins (e.g. haloalkane dehalogenase^31^) and proteins which self-assemble into more complex structures (e.g. the MS2 virus like particle^19^), it has been difficult to apply toward proteins involved in the structure and function of living systems. One significant barrier has been the need to incorporate the library into the genome to maintain the native expression and regulation of the gene. With this work, we demonstrate that methods such as λ Red recombineering can be a scalable tactic for quickly incorporating a CCM library directly into the genome. This opens the door to apply the deep mutational scanning and protein fitness landscape framework to new cellular targets, including other multimeric protein structures similar to secretion systems, and even proteins involved in other cellular functions, like signaling or regulation.

One immediate advantage of the CCM approach is to engineer systems within living cells to achieve a desired function. In this work, we discovered many variants with higher secretion scores than PrgI^WT^ and four variants that conferred an ∼50% increase in secretion titer above that observed with PrgI^WT^, even in the context of our optimal engineered system. In contrast, the alanine scan of the *S. flexneri* T3SS PrgI homolog MxiH was only able to identify 7 variants with enhanced secretion titer^15^.

The real power in the CCM approach is the information gleaned from the resulting fitness landscapes. This information allows us to generate new hypotheses to deepen our understanding of the protein structure/function relationship, which in turn can inspire new engineering approaches to further our control of the system. In this work, we not only found single mutations which increased secretion titer, but we were also able to analyze which positions were most tolerant of mutations and what kind of amino acid substitutions were allowable to maintain function. These granular data surpass what is possible through conservation studies of protein homologs, for example. By connecting the SFL with the assembled PrgI structure, we discerned that many favorable substitutions occurred in the C-terminal helix positions which make up the hydrophobic groove in the needle lumen. These positions are highly conserved, so it was not obvious from conservation alone that modifying the hydrophobic groove would not only be accepted but substantially enhance secretion. Even more surprising was a preference for large hydrophobic substitutions in high-secreting variants, a finding that would have been unlikely to be reached using rational design methods. Future analysis of the SFL and the protein structure could further our understanding of the T3SS’s assembly and how proteins are secreted through the channel.

The secretion fitness landscape of PrgI does not provide a roadmap for mutations that enhance native T3SS fitness. Rather, it might provide a roadmap for mutations that decrease native T3SS fitness. The native secretion apparatus did not evolve to secrete a maximum amount of each native substrate—secretion titers of native substrates via the T3SS are tightly controlled in its native context^32,33^. A recent alanine scan of PrgI revealed that mutations that increased secretion of native effectors produced variable invasion levels of *S. enterica* Typhimurium into human intestine epithelial cells, indicating that some mutations interrupted the native program of the T3SS^17^. The high average secretion scores of cysteine and methionine underline this difference, as the native PrgI lacks cysteine, methionine, histidine, and tryptophan.

Finally, this work highlights an underutilized approach toward engineering secretion systems. Typically, efforts have focused on rewiring regulation to maximize expression of the system^5^, modifying the secreted protein cargo to be secretable^34,35^, and porting the machinery to other organisms^24,36^. Modifications to the structural proteins of the secretion system have mostly been limited to targeted insertions or deletions^16,37^. We have shown that even though they are highly conserved, the engineering space for T3SS structural proteins is not limited. This work provides a blueprint for engineering other T3SS proteins and proteins of other secretion systems to further the goal of high titer, low cost of valuable heterologous proteins.

## Materials and Methods

### Strains and Growth Conditions for Secretion Experiments

Strains, plasmids, and primers used are listed in **Tables S1-S3**, respectively. Secretion experiments were started by growing a single colony in the lysogeny broth Lennox formulation (LB-L) (10 g/L tryptone, 5 g/L yeast extract, 5 g/L NaCl) with appropriate antibiotics (34 μg/mL chloramphenicol for P*_sicA_* vectors, 50 μg/mL kanamycin for P*_lacUV5_ hilA*) for 12-16 hours overnight in an orbital shaker at 37°C and 225 rpm unless otherwise specified. Overnight cultures were diluted 1:100 into the appropriate medium supplemented with appropriate antibiotics and 100 μg/mL isopropyl β-D-1-thiogalactopyranoside (IPTG) if the strain carried P*_lacUV5_ hilA*. All culturing steps were performed in 5 mL cultures in 24-well deepwell plates (Axygen) unless otherwise specified. Secretion was performed at 37°C and 225 rpm in an orbital shaker for eight hours unless otherwise specified. The secreted fraction was harvested by centrifuging cultures at 4000 x g for 10 minutes. Sodium dodecyl sulfate-polyacrylamide gel electrophoresis (SDS-PAGE) samples for the secretion fraction were prepared by adding supernatant to Laemmli buffer in a 3:1 ratio; SDS-PAGE samples for whole culture lysate were prepared by adding cell suspension to Laemmli buffer in a 1:2 ratio. All SDS-PAGE samples were boiled at 95°C for 5 minutes immediately after preparation.

### Protein Separation, Western Blotting, and Densitometry

Samples were separated by SDS-PAGE and transferred to a polyvinylidene fluoride membrane (PVDF, Millipore) for western blotting. If necessary, samples were further diluted in 1X Laemmli buffer such that all band signals were within twofold of the average signal across the blot. Membranes were probed with mouse anti-FLAG per manufacturer’s instructions (Sigma Aldrich). A secondary labeling step was performed with goat anti-mouse IgG (H+L) HRP conjugate according to manufacturer’s instructions (Thermo Fisher) to facilitate chemiluminescent detection. Bands were detected with the SuperSignal West Pico Plus or SuperSignal West Femto (for the P41 mutants) substrates (Thermo Fisher) and a ChemiDoc XRS+ imaging system (Bio-Rad).

All relative protein quantities from western blotting were calculated by performing densitometry using ImageJ or Image Lab software (Bio-Rad) and normalizing to the average of the replicates of the PrgI^WT^ samples. Relative protein amounts were corrected for dilution if appropriate. Error bars are standard deviation on three biological replicates unless otherwise specified (P41 mutant studies used four biological replicates).

### PCR and Cloning

Primers used in this study are listed in **Table S3**. Polymerase chain reaction (PCR) was performed with Phusion DNA polymerase for Quikchanges and constructing parts for recombineering. Saturation mutagenesis at position 41 was performed by introducing mutations to *prgI* carried on a P*_lacUV5_*-inducible plasmid with a Quikchange protocol. Mutations were confirmed by Sanger sequencing. Double-stranded DNA fragments for recombineering contained the replacement gene(s) flanked 5’ and 3’ by 40 base pairs (bp) of homology to the genetic locus at which the replacement gene(s) should be inserted. The 40 bp of homology was included in oligos and attached via PCR using the primers listed in **Table S3**. The *catG-sacB* cassette was amplified from the purified genome of *E. coli* TUC01, PrgI position 41 mutants were amplified from the appropriate P*_lacUV5_*-inducible plasmid, and *sptP-phoA-2xFLAG-6xHis* was amplified from a P*_sicA_ sicP sptP-phoA-2xFLAG-6xHis* secretion plasmid. Colony PCR was performed by diluting a colony in a 50 μL PCR reaction containing the appropriate primers and amplifying with GoTaq polymerase. Correct sequences were confirmed by Sanger sequencing.

### Strain Construction for Single Genomic Modifications

Strain modifications were generated by λ Red recombineering as described by Thomason *et al* ^25^. Briefly, a colony of ASTE13 carrying the pSIM6 plasmid was inoculated in LB-L with 30 μg/mL carbenicillin and grown at 30°C and 225 rpm for 16-20 hours. The overnight culture was diluted 1:70 into 35 mL of LB-L and grown at 30°C until OD_600_ reach 0.4-0.6. The culture was washed twice with 30 mL ice-cold sterile ddH_2_O and centrifugation at 4600 x g for 3 minutes to collect the cells. After the second wash, cells were resuspended in ∼400 μL of ice-cold sterile ddH_2_O. Aliquots of 50 μL resuspended cells were mixed with 200 ng of the appropriate PCR fragment and electroporated at 1800V for 5 milliseconds. A negative control containing no added DNA was also electroporated. Cells were mixed with 950 μL Super Optimal broth with Catabolite repression (SOC) medium immediately after electroporation and either recovered at 30°C for an hour for *cat-sacB* cassette introduction (first step of recombineering) ors transferred to a test tube containing 9 mL of LB-L and grown at 37°C, 225 rpm for four hours for *cat-sacB* removal and replacement (second step of recombineering). Cells were diluted serially to 10^-3^ in sterile phosphate buffered saline (PBS). 200 μL of diluted cells was plated on 6% sucrose agar and grown at 37°C overnight. The second step of recombineering for the *ΔinvA* knockout replaced the *cat-sacB* cassette by electroporating 1 μL of a 10 μM solution of a single 60 bp oligo containing the first and last 30 bp of the *invA* gene.

### Library Construction

A library of gene blocks carrying all possible amino acid substitutions was synthesized and pooled by Twist Biosciences. Codons were fully randomized (“NNN”, meaning any nucleotide at all three codon positions), but the library excluded wild-type residues and stop codons. Residues 1-6 and 76-80 were not modified. The lyophilized DNA from Twist Biosciences was reconstituted in ultrafiltered water to a concentration of 200 ng/μL. Recombineering was performed with an ASTE13 *sptP::sptP(1-167)-phoA-2xFLAG-6xHis prgI::cat-sacB* as described above with the following modifications. 200 ng (4 μL of a 50 ng/μL resuspended solution) of the library was transformed into 100 μL of recombination-competent cells via electroporation at 1800V and 5 milliseconds. A negative control containing no added DNA was also electroporated. Cells were immediately mixed with 900 μL SOC medium and transferred to a 14 mL disposable culture tube (Fisherbrand) containing 2 mL of LB-L for a four-hour recovery at 37°C and 225 rpm. Recombination efficiency was assessed by plating 200 μL of cells diluted serially to 10^-3^ in sterile PBS from both the library and the negative control on 6% sucrose agar and allowing colonies to develop at room temperature for 24 hours. The remainder of the culture was mixed with 60% glycerol in a 1:3 ratio and aliquoted into three cryovials for storage at -80°C. Before storage, 2 x 33 μL aliquots of the glycerol mixture were diluted to facilitate plating single colonies for screening. The first aliquot was diluted in 1.2 mL sterile PBS, and the second aliquot was diluted in 1.2 mL PBS with 15% glycerol and frozen at -80°C. The 1.2 mL aliquot without glycerol was further split into 3 x 400 μL aliquots, and each was plated on a 15 cm agar plate with 6% sucrose LB-agar. Colonies developed for 24 hours at room temperature.

### Library Screening

Single colonies were inoculated in 0.5 mL LB-L in a 2 mL square 96-well deepwell plate (Axygen) and grown overnight at 37°C, 350 rpm. ASTE13 *sptP::sptP(1-167)-phoA-2xFLAG-6xHis*, ASTE13 *sptP::sptP(1-167)-phoA-2xFLAG-6xHis prgI::catG-sacB*, and ASTE13 *sptP::sptP(1-167)-phoA-2xFLAG-6xHis ΔinvA* were included in each deepwell plate as controls. Overnight cultures were stored for analysis and high-throughput sequencing by diluting 180 μL of overnight culture with 60 μL 60% glycerol in a sterile, round-bottom 96-well plate (Corning), sealing the plate, and storing it at -80°C. To facilitate secretion, overnight cultures were diluted 1:100 into 0.5 mL Terrific Broth (TB) in a fresh 2 mL square 96-well deepwell plate and grown for 8 hours at 37°C, 350 rpm. The secretion fraction was harvested by pelleting cells in the deepwell plates at 4000 x g for 10 minutes, collecting 200 μL of the supernatant, and storing it in a sealed plate at 4°C.

### Alkaline Phosphatase Activity

Alkaline phosphatase activity was measured by monitoring p-nitrophenol phosphate (pNPP, Sigma) cleavage. A stock solution of 0.1 M pNPP prepared in 1 M Tris pH 8.0 was thawed from -20°C and diluted to 0.01 M in 1 M Tris pH 8.0. 20 μL of the secretion fraction was added to 140 μL of 1 M Tris pH 8.0, and 40 μL of the 0.01 M pNPP solution was added to each well. AP activity was measured on a BioTek Synergy HTX plate reader by monitoring absorbance at 405 nm at 37°C for one hour, taking measurements each minute.

### Sample Preparation for High Throughput Sequencing and Data Processing

Detailed methods of sample preparation, high-throughput sequencing, and subsequent data processing are available in the Supplemental Methods file.

## Acknowledgements

The authors would like to thank many current and former members of the Tullman-Ercek lab for making this study possible. Special thanks to Dr. Han Teng Wong, who did the initial quality trimming of the high throughput sequencing reads. Many thanks to Dr. Kevin Metcalf and Elias Valdivia for laying the groundwork for this project by developing the recombineering protocol and mapping the MxiH mutations to PrgI. Thank you to Sara Fernandez Dunne and the High Throughput Analysis Core at Northwestern for use of and support for the DigiLab HiGro shakers and the Tecan Fluent used for library reformatting. We would also like to thank Dr. Emily Hartman, Dr. Daniel Brauer, and Dr. Taylor Nichols for helpful discussions on processing and visualizing the fitness landscape data.

The authors gratefully acknowledge funding for this work. SAL, LAB, and DTE were supported in part by a National Science Foundation grant CBET-2043973 to DTE. LAB and NWK received support in the form of National Science Foundation Graduate Research Fellowships. BCI was supported in part by the National Institutes of Health Training Grant T32GM105538 through Northwestern University’s Chemistry of Life Processes Training Program. JSS was supported in part by the National Institutes of Health Training Grant T32GM008449 through Northwestern University’s Biotechnology Training Program. DTE and JSS were supported in part through the NSF Materials Research Science and Engineering Centers program (NSF DMR-2308691).

## Supplementary Data

**Supplementary Figure 1.**
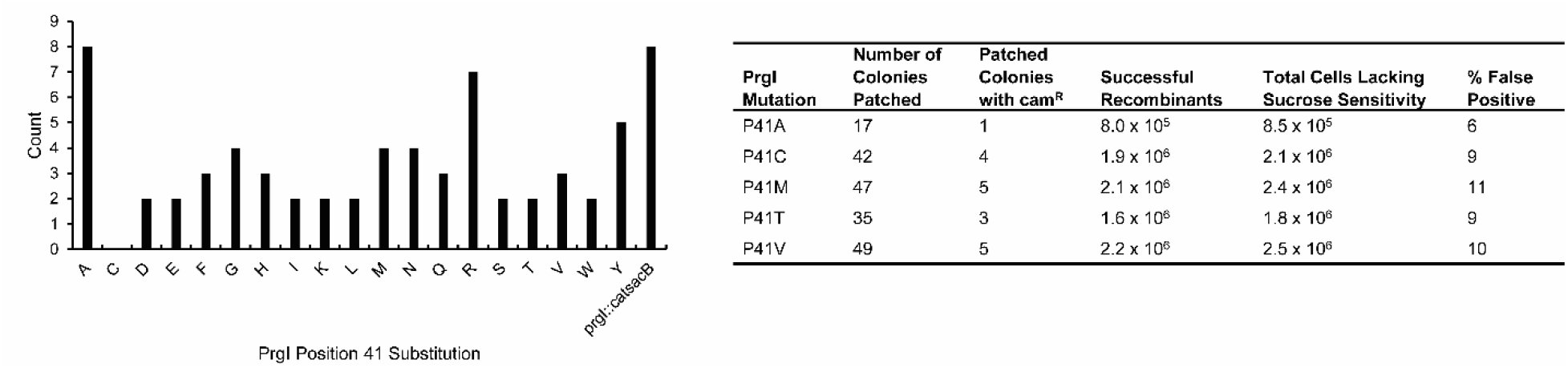
Recombineering results for the PrgI^P41^ library. An equimolar pool of all 20 fragments of the saturation mutagenesis library was introduced into an *sptP::sptP*^1–167^– *phoA prgI::catG-sacB* strain using a single λ Red recombineering event, and 68 of the resulting clones were Sanger sequenced. On the left is the count of each PrgI allele in the 68 clones. On the right are the results of conducting 5 separate recombineering events to produce individual PrgI alleles.

**Supplementary Figure 2.**
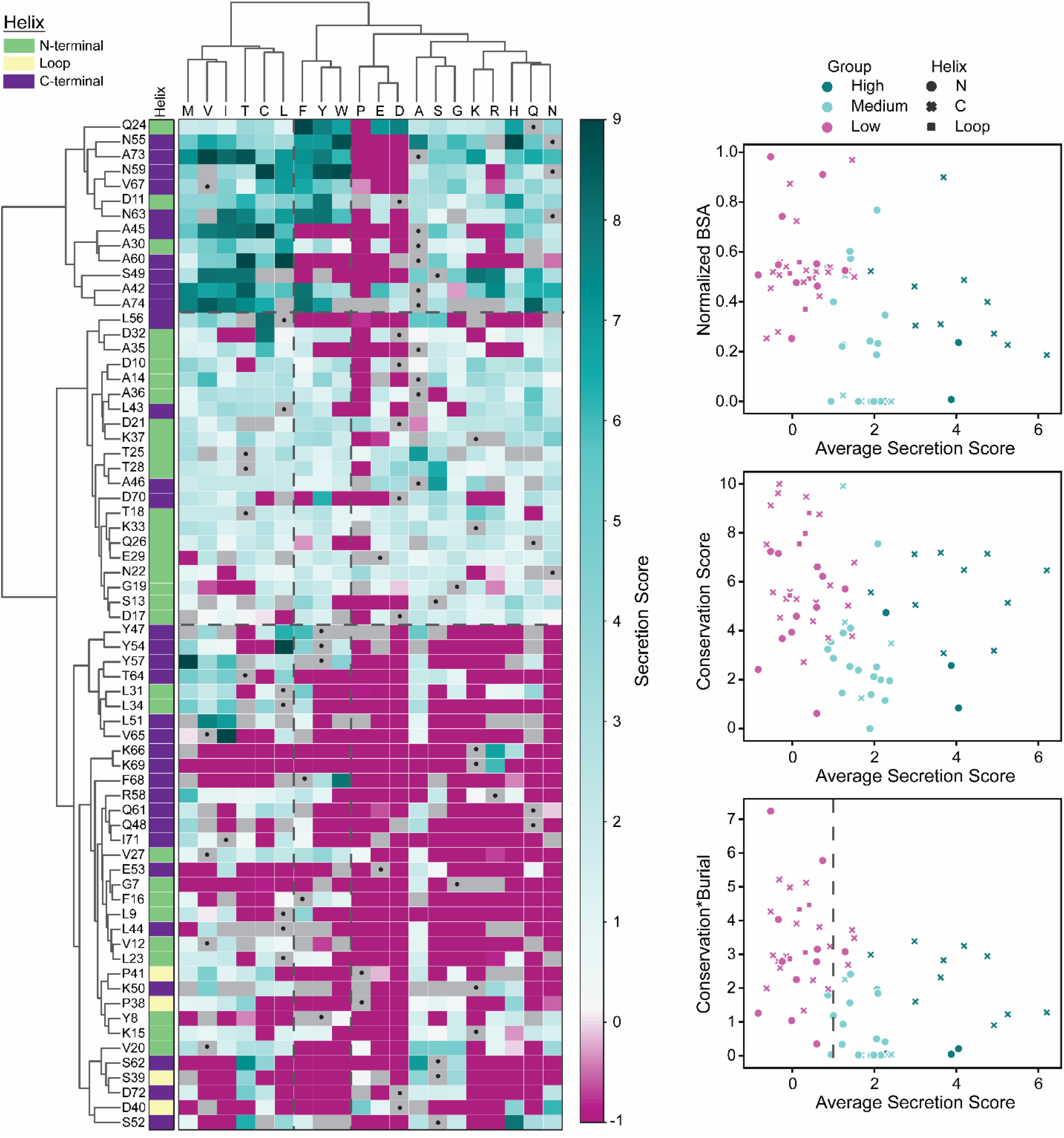
Clustering patterns of the PrgI SFL. Structural elements tended to cluster together (left), with the N-terminal helix being generally mutationally tolerant but not producing many high secreting variants. The highest secreting variants (top left cluster, dark blue) were surprisingly located in the interior N-terminal helix and were often large hydrophobic residues. Trends in the ability of BSA and conservation to predict secretion score are shown on the right.

## Supplementary Methods

### Sample preparation for high-throughput sequencing

The randomly arrayed glycerol stocks were inoculated in identical arrangements in fresh media in sterile, flat-bottom 96-well plates using a Tecan Fluent and grown with lids in DigiLab HiGro shaking stacks at 37°C, 200 rpm for 18 hours. Each clone was assigned a pool according to its relative secretion titer. A Tecan Fluent was used to reformat clones and sort them into their assigned pools. A VBA macro assigned pools according to relative secretion titer and assigned clones to a new plate and well ID to provide instructions for the Tecan Fluent. 150 μL of the fresh cell suspension was mixed with 50 μL 60% glycerol in a fresh sterile, round-bottom 96-well plate (Corning). The Tecan Fluent failed after sorting pools F-J, so the remainder were done by hand over the course of a week. Each well of the newly sorted glycerol stocks was sampled and pooled according to **Table S4** for genomic DNA purification. Genomic DNA was purified from 1 mL of each pool and 0.5 mL of the naïve library using the GenElute Bacterial Genomic DNA kit (Sigma). PCR for library preparation was conducted with Phusion polymerase. The purified genomes from the pool mixtures were amplified using the “Round 1” reaction recipe (**Table S5**) and cycling conditions (**Table S6**) with primers oLAB278 and oLAB279 (**Table S3**) to attach Illumina Nextera XT adapters. Reactions were purified using the Promega Wizard SV PCR cleanup kit. For each pool, 8 x 25 µL reactions were performed and pooled after PCR cleanup to minimize jackpot effects. A second round of PCR attached Nextera XT barcodes according to **Table S4** using the “Round 2” reaction recipe and cycling conditions in **Table S5** and **Table S6** with the pooled Round 1 reactions as templates.

### High-Throughput Sequencing Data Processing

The code for data processing using the Linux command-line interface (bash) is given following each explanation. Data were trimmed using Trimmomatic[1] with a 2-unit sliding quality window of 30 and a minimum length of 30. Sequences were cropped to 243 bp.

*java -jar trimmomatic-0.36.jar SE input_forward_HTS001.fastq.gz HTS001_trimmed SLIDINGWINDOW:2:30 MINLEN:30 CROP:243*

Reads were then aligned to the wild-type PrgI reference gene with Burrows–Wheeler Aligner (BWA-MEM)[2] and piped into Samtools[3] to convert to a bam file.

*bwa mem -p Reference/ref.fasta HTS001_trimmed.fastq | samtools view -bT Reference/ref.fasta - o HTS001.bam*

Reads that fully mapped to PrgI were kept for further analysis.

*samtools view HTS001.bam | grep “243M” | sort | less -S>HTS001.txt*

The trimmed reads were further processed to generate a secretion fitness landscape using code written in-house (see below for details).

#### Secretion Titer Score Definitions

> *m*: one of the 20 canonical amino acids.
>
> *P*_*i*_: pool of screened clones sorted by secretion titer relative to PrgI^WT^. Nonfunctional clones are in pool *A*, and pools *B* − *J* contain functional clones with relative secretion titer increasing from *B* to *J*.
>
>
> 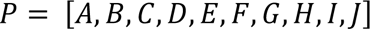
>
> *S*_*Pi*_ : a secretion titer score assigned to each pool *P*_*i*_.
>
>
> 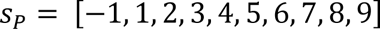
>
> *A*_*Pi,p,m*_ : an abundance score in pool *P*_*i*_, indexed by position *p* and mutation *m*.
>
> *B*_*Pi,p,m*_: a binary array recording presence (“1”) or absence (“0”) of a mutation *m* at position *p* in pool *P*_*i*_.
>
> *PA*_*Pi,p,m*_ : a percent abundance score for each mutation *m* at position *p* in pool *P*_*i*_.
>
> *B*_*Pi,p,m*_ : a secretion titer score for each mutation *m* at position *p* in pool *P*_*i*_.
>
> *BB*_*p,m*_: an array of secretion titer scores for mutation *m* at position *p*.
>
> *PAW*_*p,m*_: an array of percent abundances for mutation *m* at position *p*.
>
> *w*_*p,m*_: an array of weighted percent abundances for mutation *m* at position *p*.
>
> *WB*_*p,m*_ : a weighted average secretion titer score for each mutation *m* at position *p* in pool *P*_*i*_.
>
> 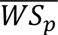: average secretion titer score per residue

#### Secretion Titer Score Calculations

Sequences were trimmed and aligned using code written in-house. Following the data processing described above, a text file was produced containing one sequencing read per line. The text file was read into Python, and only lines starting with “ATG” and ending with “TAA” were kept.

A second quality control step was implemented in Python: sequences with read counts < 20, sequences that matched the wild-type PrgI sequence, and sequences that contained more than one mutation were discarded.

Sequences that survived the more stringent quality control step were translated, and two matrices were populated: an abundance array *A* with read counts for each mutation at each position, and a binary array *B* containing a “1” for a mutation present at a position and a “0” for a mutation not present at a position.

Percent abundance for each mutation at each position was generated by dividing by the total number of read counts:

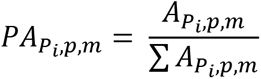

A secretion titer score was assigned for *B*_*Pi,p,m*_ = 1 to generate a new matrix for each pool with the appropriate secretion titer score at mutations that were present in that pool:

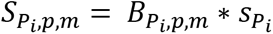

A weighted average secretion score was calculated using *PA*_*Pi,p,m*_ to compensate for the appearance of mutations in multiple pools. Secretion titer scores and percent abundances from all pools were collected into arrays for each mutation *m* at position *p*:

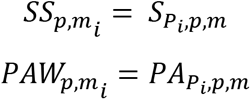

An array of weights was generated from the percent abundances and multiplied by the list of secretion scores to calculate a weighted average secretion titer score:

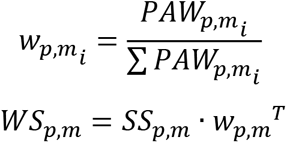

*WB*_*p,m*_ is plotted in **Fig 4**. The average secretion score per residue was calculated by averaging *WB*_*p,m*_ for each *p*, ignoring missing values:

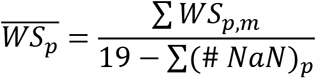

## Supplementary Tables

**Table S1.** Strains used in this study.

| Strain | Genotype | Reference |
| --- | --- | --- |
| ASTE13 | LT2-derived lab strain similar to DW01 | This study;<br>DW01 [4] |
| ASTE13 $\Delta prgI$ | $\Delta prgI$ | [5] |
| ASTE13 $prgI::catsacB$ | $prgI::catsacB$ | [6] |
| DTE509 | ASTE13 $prgI::prgI^{L9A}$ | [6] |
| DTE510 | ASTE13 $prgI::prgI^{Q48A}$ | [6] |
| DTE511 | ASTE13 $prgI::prgI^{Y54A}$ | [6] |
| DTE512 | ASTE13 $prgI::prgI^{D70A}$ | [6] |
| DTE513 | ASTE13 $prgI::prgI^{P41A}$ | [6] |
| DTE514 | ASTE13 $prgI::prgI^{P41C}$ | This study |
| DTE515 | ASTE13 $prgI::prgI^{P41D}$ | This study |
| DTE516 | ASTE13 $prgI::prgI^{P41E}$ | This study |
| DTE517 | ASTE13 $prgI::prgI^{P41F}$ | This study |
| DTE518 | ASTE13 $prgI::prgI^{P41G}$ | This study |
| DTE519 | ASTE13 $prgI::prgI^{P41H}$ | This study |
| DTE520 | ASTE13 $prgI::prgI^{P41I}$ | This study |
| DTE521 | ASTE13 $prgI::prgI^{P41K}$ | This study |
| DTE522 | ASTE13 $prgI::prgI^{P41L}$ | This study |
| DTE523 | ASTE13 $prgI::prgI^{P41M}$ | This study |
| DTE524 | ASTE13 $prgI::prgI^{P41N}$ | This study |
| DTE525 | ASTE13 $prgI::prgI^{P41Q}$ | This study |
| DTE526 | ASTE13 $prgI::prgI^{P41R}$ | This study |
| DTE527 | ASTE13 $prgI::prgI^{P41S}$ | This study |
| DTE528 | ASTE13 $prgI::prgI^{P41T}$ | This study |
| DTE529 | ASTE13 $prgI::prgI^{P41V}$ | This study |
| DTE530 | ASTE13 $prgI::prgI^{P41W}$ | This study |
| DTE531 | ASTE13 $prgI::prgI^{P41Y}$ | This study |
| sLAB190 | ASTE13 $sptP::sptP^{(1-167)}-phoA-2xFLAG-6xHis$ | This study |
| sLAB191 | ASTE13 $sptP::sptP^{(1-167)}-phoA-2xFLAG-6xHis prgI::catsacB$ | This study |
| sLAB192 | ASTE13 $sptP::sptP^{(1-167)}-phoA-2xFLAG-6xHis prgI::prgI^{P41A}$ | This study |
| sLAB193 | ASTE13 $sptP::sptP^{(1-167)}-phoA-2xFLAG-6xHis prgI::prgI^{P41C}$ | This study |
| sLAB194 | ASTE13 $sptP::sptP^{(1-167)}-phoA-2xFLAG-6xHis prgI::prgI^{P41M}$ | This study |
| sLAB195 | ASTE13 $sptP::sptP^{(1-167)}-phoA-2xFLAG-6xHis prgI::prgI^{P41T}$ | This study |
| sLAB196 | ASTE13 $sptP::sptP^{(1-167)}-phoA-2xFLAG-6xHis prgI::prgI^{P41V}$ | This study |
| sLAB203 | ASTE13 $sptP::sptP^{(1-167)}-phoA-2xFLAG-6xHis prgI::prgI^{P41D}$ | This study |
| sLAB204 | ASTE13 *sptP::sptP<sup>(1-167)</sup>-phoA-2xFLAG-6xHis prgI::prgI<sup>P41E</sup>* | This study |
| sLAB205 | ASTE13 *sptP::sptP<sup>(1-167)</sup>-phoA-2xFLAG-6xHis prgI::prgI<sup>P41F</sup>* | This study |
| sLAB206 | ASTE13 *sptP::sptP<sup>(1-167)</sup>-phoA-2xFLAG-6xHis prgI::prgI<sup>P41G</sup>* | This study |
| sLAB207 | ASTE13 *sptP::sptP<sup>(1-167)</sup>-phoA-2xFLAG-6xHis prgI::prgI<sup>P41H</sup>* | This study |
| sLAB208 | ASTE13 *sptP::sptP<sup>(1-167)</sup>-phoA-2xFLAG-6xHis prgI::prgI<sup>P41I</sup>* | This study |
| sLAB209 | ASTE13 *sptP::sptP<sup>(1-167)</sup>-phoA-2xFLAG-6xHis prgI::prgI<sup>P41K</sup>* | This study |
| sLAB210 | ASTE13 *sptP::sptP<sup>(1-167)</sup>-phoA-2xFLAG-6xHis prgI::prgI<sup>P41L</sup>* | This study |
| sLAB211 | ASTE13 *sptP::sptP<sup>(1-167)</sup>-phoA-2xFLAG-6xHis prgI::prgI<sup>P41N</sup>* | This study |
| sLAB212 | ASTE13 *sptP::sptP<sup>(1-167)</sup>-phoA-2xFLAG-6xHis prgI::prgI<sup>P41Q</sup>* | This study |
| sLAB213 | ASTE13 *sptP::sptP<sup>(1-167)</sup>-phoA-2xFLAG-6xHis prgI::prgI<sup>P41R</sup>* | This study |
| sLAB214 | ASTE13 *sptP::sptP<sup>(1-167)</sup>-phoA-2xFLAG-6xHis prgI::prgI<sup>P41S</sup>* | This study |
| sLAB215 | ASTE13 *sptP::sptP<sup>(1-167)</sup>-phoA-2xFLAG-6xHis prgI::prgI<sup>P41W</sup>* | This study |
| sLAB216 | ASTE13 *sptP::sptP<sup>(1-167)</sup>-phoA-2xFLAG-6xHis prgI::prgI<sup>P41Y</sup>* | This study |
| sLAB305 | ASTE13 *sptP::sptP<sup>(1-167)</sup>-phoA-2xFLAG-6xHis $\Delta invA$* | This study |

**Table S2.**
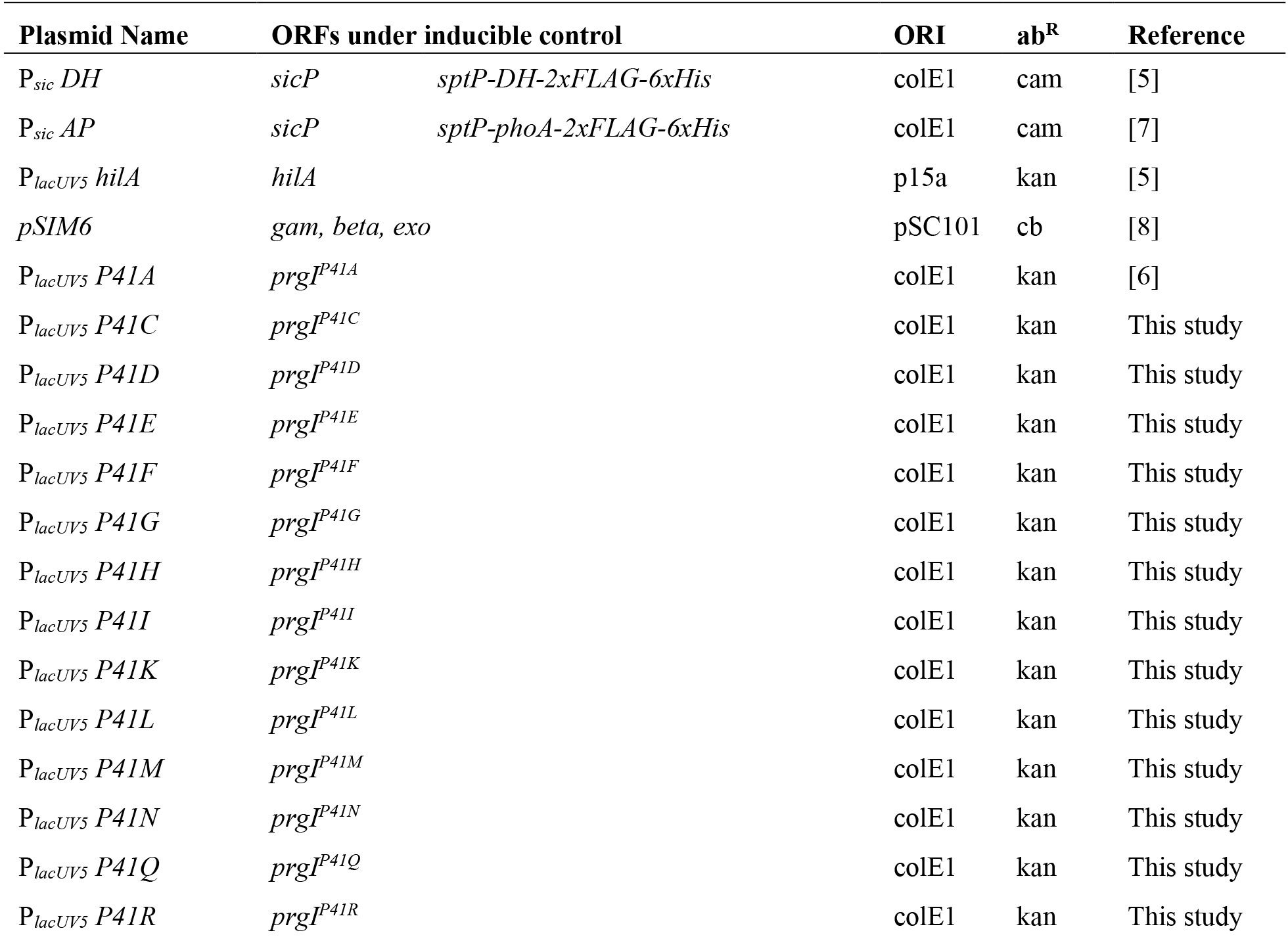
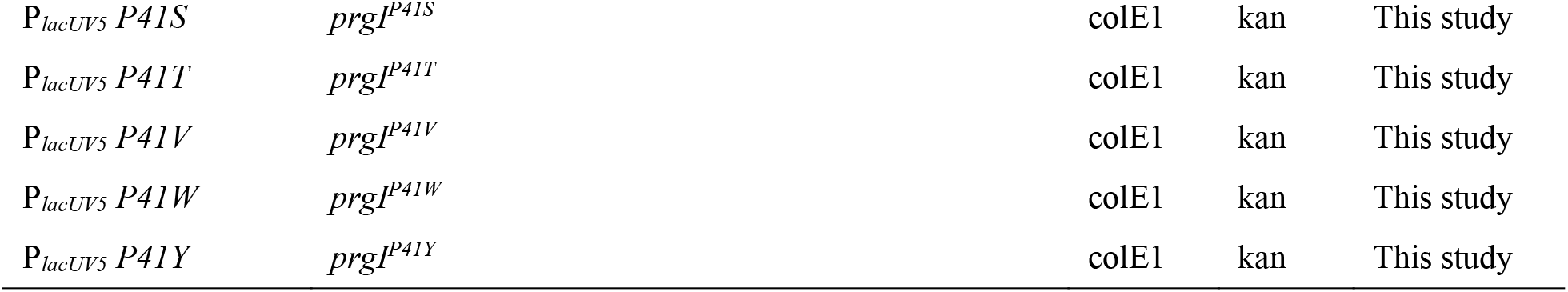
Plasmids used in this study.

**Table S3.**
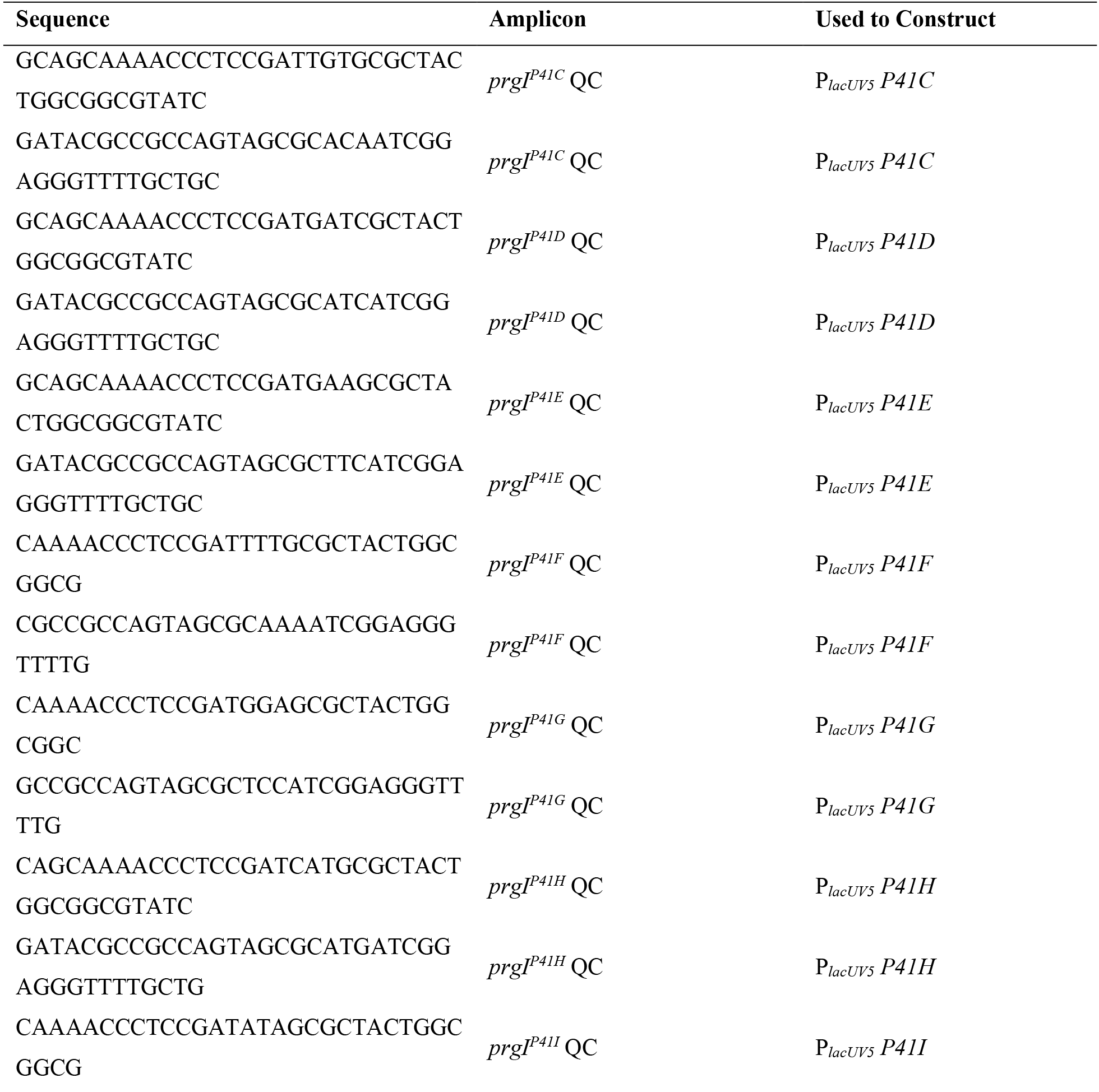
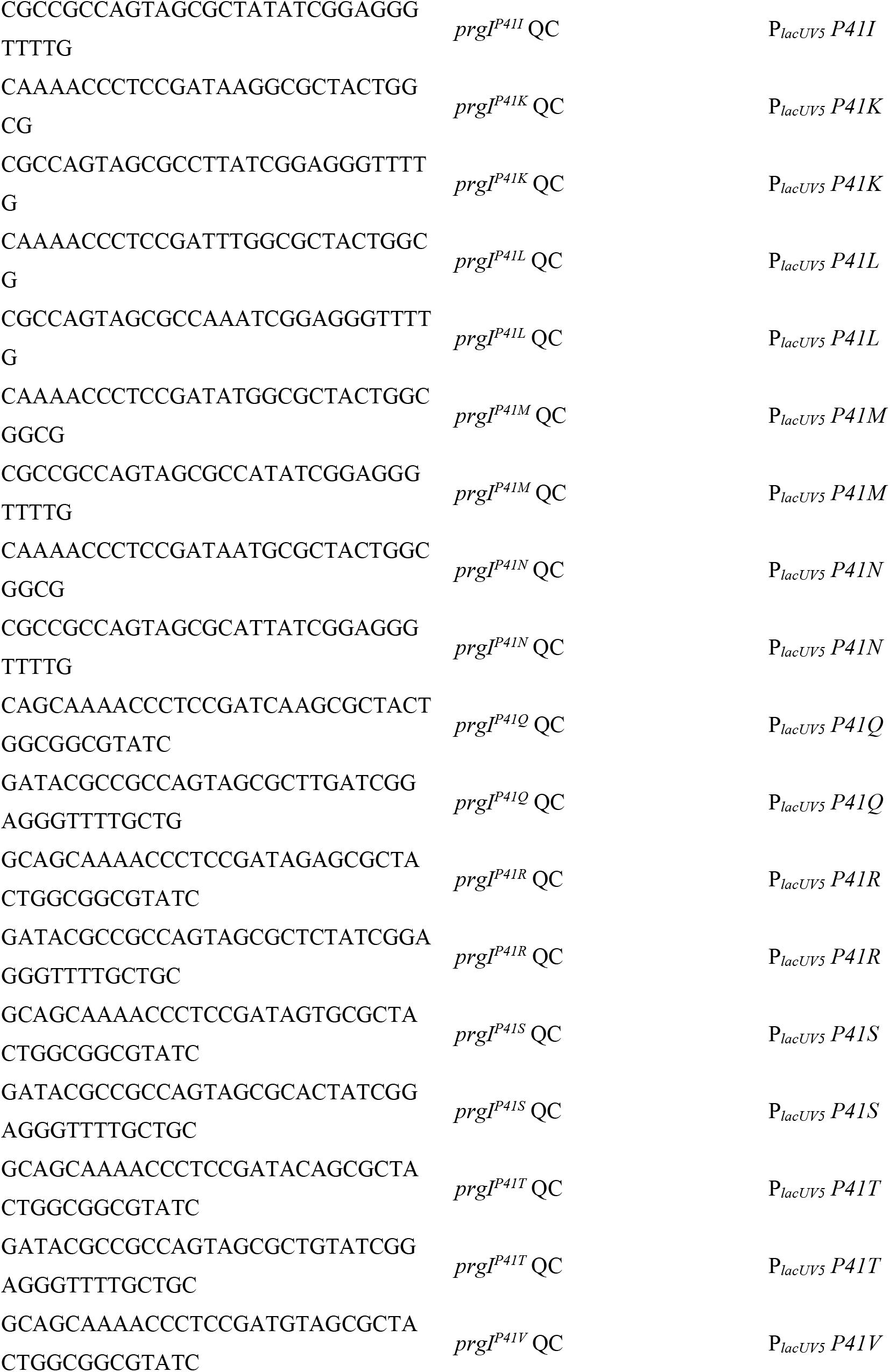
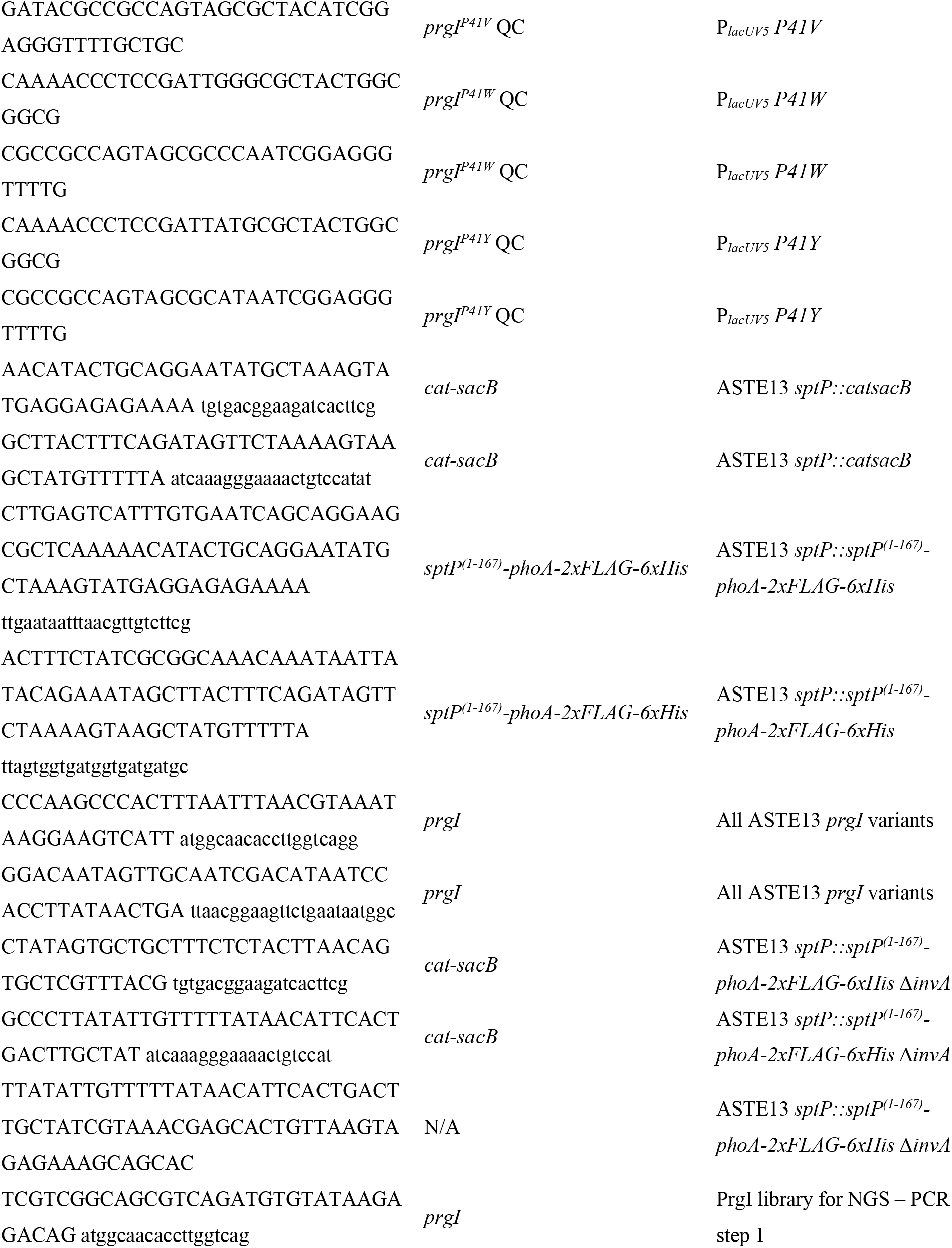
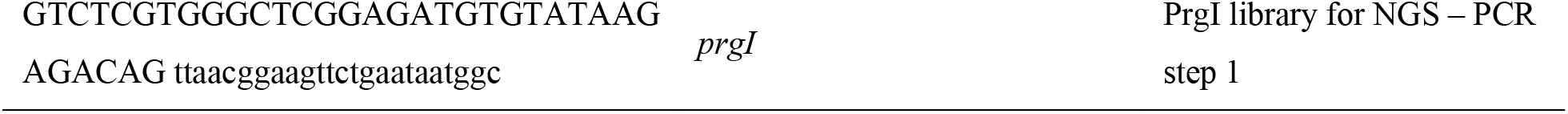
Primers used in this study.

**Table S4.** Pools for high throughput sequencing according to relative secretion titer.

| Pool | Relative Secretion<br>Titer | Number of<br>Clones | Sample Volume<br>for Mixture (μL) | Nextera XT<br>Primer i5 | Nextera XT<br>Primer i7 |
| --- | --- | --- | --- | --- | --- |
| A | 0.01-0.6 | 2015 | 5 | N707 | S502 |
| B | 0.6-0.8 | 575 | 10 | N710 | S502 |
| C | 0.8-1.0 | 691 | 10 | N711 | S502 |
| D | 1.0-1.2 | 480 | 20 | N712 | S502 |
| E | 1.2-1.4 | 236 | 20 | N714 | S502 |
| F | 1.4-1.6 | 151 | 20 | N705 | S503 |
| G | 1.6-1.8 | 77 | 30 | N706 | S503 |
| H | 1.8-2.0 | 75 | 30 | N707 | S503 |
| I | 2.0-2.5 | 70 | 30 | N710 | S503 |
| J | >2.5 | 36 | 50 | N711 | S503 |

**Table S5.** PCR reactions for high throughput sequencing library preparation.

| Component | Round 1 (25 μL x 8 per pool) | Round 2 (50 μL x 1 per pool) |
| --- | --- | --- |
| 5X HF Buffer (NEB) | 5 μL | 10 μL |
| 10 mM dNTPs (NEB) | 0.5 μL | 1 μL |
| 10 μM FWD primer | 1.25 μL | 2.5 μL |
| 10 μM REV primer | 1.25 μL | 2.5 μL |
| Template DNA | 2.5 μL of 5 ng/μL gDNA | 5 μL purified and combined Round<br>1 reaction |
| Phusion (NEB) | 0.25 μL | 0.5 μL |
| H<sub>2</sub>O | 14.25 μL | 28.5 μL |

**Table S6.** PCR cycling conditions for high throughput sequencing library preparation.

| Step | <u>Round 1 (25 μL x 8 per pool)</u> |  | <u>Round 2 (50 μL x 1 per pool)</u> |  |
| --- | --- | --- | --- | --- |
|  | T (°C) | Time (sec) | T (°C) | Time (sec) |
| Initial Denaturation | 98 | 60 | 98 | 30 |
| Amplification | 98 | 10 | 98 | 10 |
| (Round 1 – 22 cycles) | 62 | 15 | 61 | 15 |
| (Round 2 – 8 cycles) | 72 | 30 | 72 | 30 |
| Elongation | 72 | 300 | 72 | 300 |
| Hold | 4 | indefinite | 4 | indefinite |

## References

1. Derman, A. I., Prinz, W. A., Belin, D. & Beckwith, J. Mutations That Allow Disulfide Bond Formation in the Cytoplasm of Escherichia coli. Science 262, 1744–1747 (1993).

2. Dong, H., Nilsson, L. & Kurland, C. G. Gratuitous overexpression of genes in Escherichia coli leads to growth inhibition and ribosome destruction. J. Bacteriol. 177, 1497–1504 (1995).

3. Orozco-Hidalgo, M. T. et al. Engineering High-Yield Biopolymer Secretion Creates an Extracellular Protein Matrix for Living Materials. mSystems 6, 10.1128/msystems.00903-20 (2021).

4. Burdette, L. A., Leach, S. A., Wong, H. T. & Tullman-Ercek, D. Developing Gram-negative bacteria for the secretion of heterologous proteins. Microb. Cell Factories 17, 196 (2018).

5. Metcalf, K. J., Finnerty, C., Azam, A., Valdivia, E. & Tullman-Ercek, D. Using transcriptional control to increase titer of secreted heterologous proteins by the type III secretion system. Appl. Environ. Microbiol. AEM.01330–14 (2014) doi:10.1128/AEM.01330-14.

6. Masi, M. & Wandersman, C. Multiple Signals Direct the Assembly and Function of a Type 1 Secretion System. J. Bacteriol. 192, 3861–3869 (2010).

7. Guo, S., Alshamy, I., Hughes, K. T. & Chevance, F. F. V. Analysis of Factors That Affect FlgM-Dependent Type III Secretion for Protein Purification with Salmonella enterica Serovar Typhimurium. J. Bacteriol. 196, 2333–2347 (2014).

8. Azam, A., Li, C., Metcalf, K. J. & Tullman-Ercek, D. Type III secretion as a generalizable strategy for the production of full-length biopolymer-forming proteins. Biotechnol. Bioeng. 113, 2313–2320 (2016).

9. Burdette, L. A., Wong, H. T. & Tullman-Ercek, D. An optimized growth medium for increased recombinant protein secretion titer via the type III secretion system. Microb. Cell Factories 20, 44 (2021).

10. Marlovits, T. C. et al. Structural Insights into the Assembly of the Type III Secretion Needle Complex. Science 306, 1040–1042 (2004).

11. Bank, R. P. D. RCSB PDB - 6DWB: Structure of the Salmonella SPI-1 type III secretion injectisome needle filament. https://www.rcsb.org/structure/6DWB.

12. Büttner, D. Protein Export According to Schedule: Architecture, Assembly, and Regulation of Type III Secretion Systems from Plant- and Animal-Pathogenic Bacteria. Microbiol. Mol. Biol. Rev. (2012) doi:10.1128/MMBR.05017-11.

13. Kubori, T. et al. Supramolecular Structure of the Salmonella typhimurium Type III Protein Secretion System. Science 280, 602–605 (1998).

14. Davis, A. J. & Mecsas, J. Mutations in the Yersinia pseudotuberculosis Type III Secretion System Needle Protein, YscF, That Specifically Abrogate Effector Translocation into Host Cells. J. Bacteriol. 189, 83–97 (2007).

15. Kenjale, R. et al. The needle component of the type III secreton of Shigella regulates the activity of the secretion apparatus. J. Biol. Chem. 280, 42929–42937 (2005).

16. Glasgow, A. A., Wong, H. T. & Tullman-Ercek, D. A Secretion-Amplification Role for Salmonella enterica Translocon Protein SipD. ACS Synth. Biol. 6, 1006–1015 (2017).

17. Guo, E. Z. et al. A polymorphic helix of a Salmonella needle protein relays signals defining distinct steps in type III secretion. PLOS Biol. 17, e3000351 (2019).

18. Siloto, R. M. P. & Weselake, R. J. Site saturation mutagenesis: Methods and applications in protein engineering. Biocatal. Agric. Biotechnol. 1, 181–189 (2012).

19. Hartman, E. C. et al. Quantitative characterization of all single amino acid variants of a viral capsid-based drug delivery vehicle. Nat. Commun. 9, 1385 (2018).

20. Darboe, N., Kenjale, R., Picking, W. L., Picking, W. D. & Middaugh, C. R. Physical characterization of MxiH and PrgI, the needle component of the type III secretion apparatus from Shigella and Salmonella. Protein Sci. Publ. Protein Soc. 15, 543–552 (2006).

21. Metcalf, K. J. *Engineering Heterologous Protein Secretion for Improved Production*. (University of California, Berkeley, 2016).

22. Chatterjee, S., Chaudhury, S., McShan, A. C., Kaur, K. & De Guzman, R. N. Stucture and biophysics of Type III secretion in bacteria. Biochemistry (2013) doi:10.1021/bi400160a.

23. Kimbrough, T. G. & Miller, S. I. Contribution of Salmonella typhimurium type III secretion components to needle complex formation. Proc. Natl. Acad. Sci. U. S. A. 97, 11008–11013 (2000).

24. Song, M. et al. Control of type III protein secretion using a minimal genetic system. Nat. Commun. 8, 14737 (2017).

25. Thomason, L. et al. Recombineering: Genetic Engineering in Bacteria Using Homologous Recombination. in *Current Protocols in Molecular Biology* (John Wiley & Sons, Inc., 2001).

26. Widmaier, D. M. et al. Engineering the Salmonella type III secretion system to export spider silk monomers. Mol. Syst. Biol. 5, 309 (2009).

27. Metcalf, K. J. et al. Proteins adopt functionally active conformations after type III secretion. Microb. Cell Factories 15, 213 (2016).

28. Pettersen, E. F. et al. UCSF Chimera--a visualization system for exploratory research and analysis. J. Comput. Chem. 25, 1605–1612 (2004).

29. Loquet, A. et al. Atomic model of the type III secretion system needle. Nature (2012) doi:10.1038/nature11079.

30. Hu, J. et al. Cryo-EM analysis of the T3S injectisome reveals the structure of the needle and open secretin. Nat. Commun. 9, 3840 (2018).

31. Gray, K. A. et al. Rapid Evolution of Reversible Denaturation and Elevated Melting Temperature in a Microbial Haloalkane Dehalogenase. Adv. Synth. Catal. 343, 607–617 (2001).

32. Schlumberger, M. C. et al. Real-time imaging of type III secretion: Salmonella SipA injection into host cells. Proc. Natl. Acad. Sci. U. S. A. 102, 12548–12553 (2005).

33. Galán, J. E., Lara-Tejero, M., Marlovits, T. C. & Wagner, S. Bacterial Type III Secretion Systems: Specialized Nanomachines for Protein Delivery into Target Cells. Annu. Rev. Microbiol. 68, 415–438 (2014).

34. Bakkes, P. J., Jenewein, S., Smits, S. H. J., Holland, I. B. & Schmitt, L. The Rate of Folding Dictates Substrate Secretion by the Escherichia coli Hemolysin Type 1 Secretion System. J. Biol. Chem. 285, 40573–40580 (2010).

35. Schwarz, C. K. W., Landsberg, C. D., Lenders, M. H. H., Smits, S. H. J. & Schmitt, L. Using an E. coli Type 1 secretion system to secrete the mammalian, intracellular protein IFABP in its active form. J. Biotechnol. 159, 155–161 (2012).

36. González-Prieto, C. & Lesser, C. F. Rationale redesign of type III secretion systems: toward the development of non-pathogenic E. coli for in vivo delivery of therapeutic payloads. Curr. Opin. Microbiol. 41, 1–7 (2018).

37. Rüssmann, H., Kubori, T., Sauer, J. & Galán, J. E. Molecular and functional analysis of the type III secretion signal of the Salmonella enterica InvJ protein. Mol. Microbiol. 46, 769–779 (2002).

## Supplementary references

[1] Bolger, A. M., Lohse, M. & Usadel, B. Trimmomatic: a flexible trimmer for Illumina sequence data. Bioinformatics 30, 2114–2120 (2014).

[2] Li, H. & Durbin, R. Fast and accurate short read alignment with Burrows-Wheeler transform. Bioinformatics 25, 1754–1760 (2009).

[3] Li, H. et al. The sequence alignment/map format and SAMtools. Bioinformatics 25, 2078–2079 (2009).

[4] Song M, Sukovich DJ, Ciccarelli L, Mayr J, Fernandez-Rodriguez J, Mirsky EA, et al. Control of type III protein secretion using a minimal genetic system. Nature Communications. 2017;8:14737.

[5] Metcalf KJ, Finnerty C, Azam A, Valdivia E, Tullman-Ercek D. Using Transcriptional Control To Increase Titers of Secreted Heterologous Proteins by the Type III Secretion System. Applied and Environmental Microbiology. 2014;80:5927–34.

[6] Metcalf KJ. Engineering heterologous protein secretion for improved production. University of California, Berkeley; 2016.

[7] Metcalf KJ, Bevington JL, Rosales SL, Burdette LA, Valdivia E, Tullman-Ercek D. Proteins adopt functionally active conformations after type III secretion. Microbial Cell Factories. 2016;15:213.

[8] Thomason LC, Sawitzke JA, Li X, Costantino N, Court DL. Recombineering: Genetic Engineering in Bacteria Using Homologous Recombination. Current Protocols in Molecular Biology. 2014;106:1.16.1–1.16.39.

